# BRG1 targeting overcomes ABCC-based multidrug resistance induced by paclitaxel

**DOI:** 10.1101/2025.05.01.651609

**Authors:** Karolina Gronkowska, Sylwia Michlewska, Tomasz Płoszaj, Magdalena Strachowska, Adrianna Stępień, Maciej Borowiec, Andrzej Bednarek, Agnieszka Robaszkiewicz

## Abstract

Multidrug resistance of cancer cells is attributed to drug-induced alteration of numerous intracellular processes. Using clinically relevant models of triple-negative breast and non-small lung cancer cells we previously showed that these cells respond to repeated paclitaxel exposure by inter alia lysosome enrichment in ABCC3, ABCC5 and ABCC10, which contribute to drug sequestration in these organelles and reduced drug cytotoxicity. In this study we provide experimental evidence that transcription of above mentioned ABCC genes is enabled by BRG1-based SWI/SNF chromatin remodeling complex. Pharmacological inhibition of SWI/SNF with PFI3 or ACBI1, the PROTAC degrader of SMARCA2/4, substantially decline transcription of ABCC3, ABCC5 and ABCC10. Similar effect is caused by transient silencing of SMARCA4 (BRG1), but not SMARCA2 (BRM). The deficiency of BRG1 led to extralysosomal distribution of anticancer drugs, their deeper penetration of spheroids and substantial increase in drug cytotoxicity. Interestingly, in BRG1 deficient cell line paclitaxel triggered mutations, which reverted BRG1 truncating deletion in SMARCA4, thereby restoring SWI/SNF ATPase expression in paclitaxel-resistant cells and increasing transcription of ABCC. Acquisition of drug resistance was associated with BRG1 redistribution in the genome, de novo occurrence at the promoters of genes functionally linked to endo-lysosomal system and stronger co-occurrence with EP300. Our study indicates possible target - SWI/SNF complex for anticancer combinatorial interventions in paclitaxel-induced multidrug resistant phenotypes.

## 1. Introduction

Taxanes are successful in treating a variety of cancers due to a combination of peculiarities, in particular their unique ability to modulate various cellular processes associated with tumor growth and metabolism (apoptosis, angiogenesis, oxidative stress, etc.). This efficacy has been confirmed in the treatment of both solid and disseminated tumors. Three different taxanes are available on the market: paclitaxel (PTX), marketed under the tradenames Taxol®, Anzatax® or Paxene®, the synthetic derivative of PTX - docetaxel (DTX) (Taxotere®) and a recently synthesized cabazitaxel (CTX) (Jevtana®) [1]. PTX is a major first-line drug for the treatment of advanced non-small cell lung, breast and ovarian cancers [2–4]. At high concentrations, PTX prevents cell division by promoting an assembly of stable microtubules especially from β-tubulin heterodimers, and inhibits their depolymerization. The exposed cells are arrested in the G2/M-phase of the cell cycle and undergo apoptosis. At low concentrations, apoptosis is induced at G0 and G1/S phase either via Raf-1 kinase activation or p53/p21 depending on the administered dose [2]. However, the development of resistance to paclitaxel leads to treatment failure, tumor recurrence and limits the use of this drug. Major mechanisms of resistance include altered apoptotic pathways, overexpression of pumps that actively transport the drug out of the cells, reduced drug uptake, increased drug metabolism, alteration in the tubulin dynamic, mutations in the β-tubulin gene or expression of β-tubulin isotypes [2,5,6]. Many of these mechanisms occur at the genomic level and involve modification in the signaling routes, which then modulate activity of the subset of transcription factors and the epigenetic transformation of chromatin. Several reports have suggested that DNA methylation and epigenetic silencing of *inter alia* pro-apoptotic genes and tumor suppressors result in resistance acquisition [7].

Acquired paclitaxel resistance *via* the alteration of the cell transcriptome can involve the overexpression of ATP-binding cassette (ABC) transporters that use energy from ATP hydrolysis to translocate substances across the cell membranes. Acquisition of paclitaxel resistance has been linked to the overexpression of ABCB1 in lung cancer cells [8–10], triple negative breast cancer cells [11], ovarian cancer cells [12], ABCC5 in nasopharyngeal carcinoma cells [13] and ABCC10 in lung cancer cells [9]. In our previous study, it was shown that regardless of the alteration in gene expression, paclitaxel induced intracellular redistribution of ABCC3, ABCC5 and ABCC10 and their enrichment in lysosomes. The use of ABCC inhibitors and the transient silencing of these 3 genes considerably limited the accumulation of doxorubicin and Paclitaxel OregonGreen488 in lysosomes and sensitized cancer cells to various structurally unrelated chemotherapeutics [14]. Since these 3 transporters are mostly located in lysosomes in the tested paclitaxel-resistant cell lines, their transcriptional repression could overcome lysosome-mediated drug resistance.

Previous studies on the regulation of ABC gene transcription describe the role of transcription factors, epigenetic enzymes and chromatin remodeling in cell adaptations to paclitaxel [13,15–19]. GCN5 acetylatransferase increased the transcription of ABCB1 and ABCG2 PTX-resistant ovarian cancer cells [20], whereas EP300 was responsible for the overexpression of several ABC transporters in PTX-doxorubicin- and cisplatin-resistant breast and lung cancers [21]. The transcription activating role of the latter enzyme was found to be supported by the BRG1-SWI/SNF complex at the subset of E2F-dependent gene promoters in proliferating cells [22,23]. Expression of several ABC transporters such as ABCB1, ABCC2, ABCC11, ABCG1 and ABCG2 in breast cancer cells has been attributed to the activity of BRG1 that was also found at gene promoters of some ABC genes in MDA-MB-231 cells [24]. BRG1 and BRM are the crucial ATP-dependent and enzymatically active components of the chromatin remodeling complex SWI/SNF, they are mutually exclusive with additional subunits combinatorically incorporated to the complex in a context-dependent manner. The diversity of the composition of the remaining subunits enables highly specific interactions of the complexes with activators and repressors of transcription and the recognition of histone modifications, which in turn determine the location of the complexes within the genome and the direction of local chromatin remodeling -i.e. its tightening or loosening. Recent reports indicate that approximately 20% of human cancers contain mutations in the subunits of the SWI/SNF complex [25]. BRG1, encoded by the *SMARCA4* gene, is one of the most frequently altered elements of the complex in cancer cells. Elevated expression of BRG1 occurred in 35% - almost 100% of analyzed primary tumors and was linked with a high proliferation rate. A BRG1-dependent increase in cancer cell division was assigned to the upregulation of the expression of enzymes responsible for fatty acid and lipid biosynthesis, induction of ABC transporter expression and high transcription rate of genes encoding direct drivers of cell proliferation (signaling cascades, checkpoints and DNA replication genes) [23].

Knowing that BRG1 augments transcription of some ABC proteins in response to single doses of various drugs and that this transcription co-regulator occurs at the gene promoters of some ABC genes in non-resistant cancer cells, we tested the possible contribution of this ATPase to ABC-driven paclitaxel resistance in clinically-relevant models of paclitaxel-resistant triple-negative breast cancer (MDA-MB-231) and non-small cell lung cancer (A549). Particular attention was paid to the role of BRG1 in the regulation of *ABCC3*, *ABCC5* and *ABCC10* gene transcription due to the involvement in lysosomal drug sequestration and hence, cell protection from some anticancer drugs as was previously described in Gronkowska *et al* [14]. Finally, we tested the selective BRG1/BRM inhibitor PFI3 as well as novel SMARCA2/4 PROTAC degrader - ACBI1 as a potential sensitizer of multidrug resistant cells to standard chemotherapy drugs.

## 2. Materials and Methods

### 2.1 Materials

Non-small-cell lung cancer cell line A549 and breast cancer cell line MCF7 were purchased from ATCC. Breast cancer cell line MDA-MB-231 was purchased from Sigma Aldrich. DMEM High Glucose w/ L-Glutamine w/ Sodium Pyruvate, fetal bovine serum and antibiotics (penicillin and streptomycin) were from Biowest (CytoGen, Zgierz, Poland). KAPA SYBR® FAST Universal (2x), KAPA HiFi HotStart ReadyMix (2x), oligonucleotides for Real-time PCR, resazurin sodium salt, doxorubicin hydrochloride, daunorubicin hydrochloride, cisplatin, paclitaxel, etoposide, were from Sigma Aldrich (Poznan, Poland). PFI-3 (Cayman Chemical), ACBI1 (Cayman Chemical) and Nunc™ Lab-Tek™ Chamber Slide were ordered in Biokom (Janki/Warsaw, Poland). siRNA Control (sc-37007) and anti-MRP5 (E-10) (sc-376965) antibodies was purchased from Santa Cruz Biotechnology (AMX, Lodz, Poland). SMARCA4 Silencer Select siRNA (s13141), SMARCA2 Silencer Select siRNA (s536647), Lipofectamine RNAiMAX, OptiMem, Dynabeads™ Protein G, UltraPure™ Phenol:Chloroform:Isoamyl Alcohol (25:24:1, v/v) (#15593031), TRI Reagent™, High-Capacity cDNA Reverse Transcription Kit, BigDye® Terminator v3.1 Cycle Sequencing Kit, Hi-Di formamide, SuperSignal™ West Pico Chemiluminescent Substrate, PageRuler™ Prestained Protein Ladder (#01154870), Pierce™ Protease Inhibitor Tablets (EDTA-free; PIC), Paclitaxel Oregon Green™ 488 conjugate (Flutax-2), Lysotracker™ Deep Red, SlowFade™ Glass Soft-set Antifade Mountant (with DAPI), anti-MRP3 (ABCC3) Polyclonal Antibody (PA5101482), anti-MRP10 (ABCC10) Polyclonal Antibody (PA5101678), Goat anti-Rabbit IgG (H+L) Cross-Adsorbed Secondary Antibody, Alexa Fluor™ 546 (#A-11010), PowerUp™ SYBR® Green Master Mix, TaqMan™ Universal Master Mix II, TaqMan™ Gene Expression Assays (FAM-MGB/20X) for ACTB (Hs01064292_g1), GAPDH (Hs02786624_g1), HPRT1 (Hs03929096_g1), ABCB1 (Hs00184500_m1), ABCC1 (Hs01561483_m1), ABCC2 (Hs00960489_m1), ABCC3 (Hs00978452_m1), ABCC4 (Hs00988721_m1), ABCC5 (Hs00981089_m1), ABCC10 (Hs01056200_m1), ABCG2 (Hs01053790_m1) were from Thermofisher Scientific (Thermofisher Scientific, Warsaw, Poland). Anti - Cleaved Caspase-3 (Asp175) (5A1E) Rabbit mAb (# 9664), anti-BRG1 (D1Q7F) Rabbit mAb (#49360), anti-Histone H3 (1B1B2) Mouse mAb (#14269), anti-rabbit IgG, HRP-linked Antibody (#7074), Anti-mouse IgG, HRP-linked Antibody (#7076), Anti-rabbit IgG (H+L), F(ab’)2 Fragment (Alexa Fluor® 488 Conjugate) (#4412), Anti-mouse IgG (H+L), F(ab’)2 Fragment (PE Conjugate) (#59997S), Anti-rabbit IgG Fab2 Fragment Alexa Fluor® 594 Probes (#8889S) were from Cell Signaling Technologies (LabJOT, Warsaw, Poland). NEBNext® Ultra™ II DNA Library Prep with Sample Purification Beads (#E7104), NEBNext® Poly(A) mRNA Magnetic Isolation Module (#E7490), NEBNext® Ultra™ II RNA Library Prep Kit for Illumina® (#E7770), NEBNext® Ultra™ II DNA Library Prep with Sample Purification Beads (#E7103) and NEBNext® Multiplex Oligos for Illumina® (Index Primers Set 3) (#E7710) were from New England Biolabs (LabJOT, Warsaw, Poland). BIOFLOAT FLEX coating solution was purchased from FaCellitate (faCellitate.com). Annexin V Apoptosis Detection Kit with Propidium iodide was purchased from BioLegend (BioCourse.pl, Katowice, Poland). MGIEasy PCR-Free DNA Library Prep Set and DNBSEQ-G400RS High-throughput Sequencing Kit (FCL SE100) were from Perlan Technologies (Perlan Technologies Poland, Warsaw, Poland)

### 2.2 Cell Culture and Treatment

A549 cells were cultured in DMEM supplemented with 10% FBS and penicillin/streptomycin (50 U/ml and 50 µg/ml, respectively) in 5% CO2. Initially, MDA-MB-231 cells were cultured in F15 medium supplemented with 15% FBS and penicillin/streptomycin (50 U/mK and 50 µg/ml, respectively) without CO2 equilibration. After 5 passages, the cells were adapted to grow in DMEM supplemented with 10% FBS and penicillin/streptomycin (50 U/ml and 50 µg/ml, respectively) in 5% CO2. Paclitaxel resistance induction and characteristics of resistant cell lines was described previously [14,21].

PFI-3 (2.5 µM) and ACBI1 (0.5 µM) were added to cells 72 h before analysis or treatment with anticancer drugs. Depending on the tested parameters anticancer drugs were administrated to cells for 24 or 48h.

### 2.3 Formation of cell spheroids

Nunc™Lab-Tek™chamber slides were coated with faCellitate BIOFLOAT FLEX coating solution according to the manufacturer protocol. After 30 minutes of air-drying of the chambers, within the laminar flow hood, the cells were seeded per well at a density of 20,000 cells. Cells grow in DMEM supplemented with 10% FBS and penicillin/streptomycin (50 U/ml and 50 µg/ml, respectively) in 5% CO2 to allow spheroids formation for 21 days.

### 2.4 Transient Gene Silencing

Cells seeded per well at a density of 100,000 cells on the 24-well plate, 10,000 cells on Nunc™ Lab-Tek™ Chamber Slide or three-week spheroids were transfected using siRNA-RNAiMAX complexes according to the previously described protocol [26]. After 6 h incubation with the complexes, DMEM supplemented with 10% FBS and antibiotics was added to the desired volume and cells were grown for another 48 h to obtain transient gene silencing.

### 2.5 Real-Time PCR

For mRNA expression evaluation, total RNA was extracted from cells using TRI Reagent™. Afterwards, mRNA was reverse transcribed with the High-Capacity cDNA Reverse Transcription Kit. The expression of selected genes was measured in Bio-Rad CFX96 C1000 Touch Real-Time system, using:

1. TaqMan™gene expression assays and the TaqMan™ universal master mix II, according to the protocol provided by the manufacturer;
2. PowerTrack™ SYBR™ Green Master Mix and the manually designed primer pairs: (GAPDH Forward: 5’ TTCTTTTGCGTCGCCAGCCGA 3’, Reverse 5’

GTGACCAGGCGCCCAATACGA 3’, ACTB Forward: 5’ TGGCACCCAGCACAATGAA 3’, Reverse 5’ CTAAGTCATAGTCCGCCTAGAAGCA 3’, HPRT1 Forward: 5’ TGACACTGGCAAAACAATGCA 3’, Reverse: 5’: GGTCCTTTTCACCAGCAAGCT 3’ BRG1 Forward: 5’ AAGAAGACTGAGCCCCGACATTC 3’, Reverse 5’ CCGTTACTGCTAAGGCCTATGC 3’, ABCC3 Forward: 5 TCCTTTGCCAACTTTCTCTGCAACTAT 3’, Reverse: 5’: CTGGATCATTGTCTGTCAGATCCGT 3’, ABCC5 Forward: 5’ AGAGGTGACCTTTGAGAACGCA 3’, Reverse: 5’: CTCCAGATAACTCCACCAGACGG 3’, ABCC10 Forward: 5’ CGGGTTAAGCTTGTGACAGAGC 3’, Reverse: 5’: AACACCTTGGTGGCAGTGAGCT 3’) according to the protocol provided by the manufacturer.

mRNA level of particular genes was first normalized to housekeeping genes. The ratio between the studied and housekeeping genes was assumed to be 1 for control cells.

### 2.6 Sanger Sequencing

Total RNA was extracted from cells using TRI Reagent™. Afterwards, mRNA was reverse transcribed with the High-Capacity cDNA Reverse Transcription Kit. The fragment of BRG1 exon 15 was amplified in gradient thermocycler LifeECO (BIOER) using KAPA HiFi HotStart ReadyMix (2x) and the manually designed primer pair (Forward: 5’ CCGGGGTATGAAGTAGCTCC 3’, Reverse 5’ GTCCACCTCAGAGACGTCAT 3’). Subsequently, cDNA sequencing was performed using the BigDye® Terminator v3.1 Cycle Sequencing Kit according to the previously described procedure [27]. Sequences were analyzed using FinchTV and MEGA11 software.

### 2.7 RNA sequencing and analysis

Total RNA was isolated using TRI Reagent™. Next, 10 ng of DNA-free total RNA was used to isolate mRNA using NEBNext® Poly(A) mRNA Magnetic Isolation Module. The purified mRNA was used for RNAseq library preparation using NEBNext® Ultra™ II RNA Library Prep Kit for Illumina® (#E7770). Sample indexing was prepared using NEBNext® Multiplex Oligos for Illumina® (Index Primers Set 3) (#E7710). All of the steps were prepared according to the protocols provided by New England Biolabs. Sequencing was performed using the Illumina NextSeq 550 System. Bioinformatical analysis of the obtained data was conducted using UseGalaxy.org (Galaxy version 24.0.rc1), an open platform as previously described [28,29]. Venn diagram was created in https://bioinformatics.psb.ugent.be/webtools/Venn/ from gene lists. Annotation of downregulated genes to gene ontology was carried out in PANTHER version 17.0.

### 2.8 Chromatin immunoprecipitation, library preparation and sequencing

Chromatin immunoprecipitation was carried out according to the protocol previously described [27]. DNA was extracted using phenol:chloroform:isoamyl alcohol (25:24:1). BRG1 was immunoprecipitated with anti-BRG1 (D1Q7F) Rabbit mAb (#49360) and EP300 with p300 (D2X6N) Rabbit mAb in non-resistant and paclitaxel-resistant MDA-MB-231 cells, whereas H3K4me3 with Tri-Methyl-Histone H3 (Lys4) (C42D8) Rabbit mAb in paclitaxel-resistant MDA-MB-231 cells only. Data for tri-methylation of H3 in MDA-MB-231 were taken from Short Reads Achieve (SRA, NCBI) and the following datasets were taken for analysis: SRR4346773 and SRR4346774 (GSM2337951) [30].

BRG1 binding motifs at the selected gene promoters were amplified using KAPA SYBR® FAST Universal Master Mix, 0.1% DMSO and the following primer pairs: *ABCB1* Forward 5’ CCAATCAGCCTCACCACAGA 3’, Reverse 5’ GATTCAGCTGATGCGCGTTT 3’; *ABCC5* Forward: 5’ CTTCCGGGTTCAAGCAGTTC 3’, Reverse: 5’ AAAATACGGCGGGGTGAGG 3’, *ABCC10* Forward: 5’ TACCCTTGGTACCGCGAGA 3’, Reverse: 5’ GTAACAGGCACTGAGCACGG 3’. As a control manually designed fragments of promoters do not containing BRG1 binding motifs was used: *ABCC2* Forward 5’ AGGTCAAGGCTGCAATGAAT 3’, Reverse: 5’ CTGTCATCGACCCAACCTTT 3’. Data were normalized to samples containing non-specific IgG. 1μg of immunoprecipitated DNA fragments was converted into library for sequencing using NEBNext® Ultra™ DNA Library Prep Kit with Sample Purification Beads for Illumina® and NEBNext® Multiplex Oligos for Illumina® (Index Primers Set 3) for BRG1 and MGIEasy PCR-Free DNA Library Prep Set for EP300 and H3K4me3 according to instruction provided by manufacturers. DNA library was sequenced on NextSeq 550 or DNBSEQ-G400 in the Department of Clinical Genetics, Medical University of Lodz, and data were released as fastq files.

### 2.9 Bioinformatic analysis of ChIP-Seq data

Adapters were trimmed during FASTQ generation as an option of Illumina pipeline in BaseSpace. All other analyses were performed in Galaxy.org (Galaxy version 24.0.rc1) [29]. The read quality was checked with FastQC and reads with Qphred <30 were removed with Trimmomatic (AVGQUAL:30). Reads were mapped with BWA against human genome (hg19). Mapped reads were normalized to counts per million (-bamCoverage). Peaks were called with MACS2 at p < 0.001. BRG1 and EP300 distribution at H3K4me3 positive TSS (bedtools Intersect intervals; entry: H3K4me3 peaks and TSS 1 bp; required overlap: 1 bp) were generated by -computeMatrix (Score files: -bigwigfiles; output options: reference point, center of regions, --sortUsing mean, --averageTypeBins mean, --missingDataAsZero, -- skipZeros), then plotHeatmap with hierarchical clustering. The same method was used for analyzing distribution of EP300 at the BRG1 peak summits. Spearman correlation plot was called –multiBamSummary (by default) and –plotCorrelation (--removeOutliers).

Blacklist regions for hg19 were downloaded from ENCODE (ENCSR636HFF) [31].

### 2.10 Western Blot

For protein expression evaluation cells were lysed in RIPA buffer (supplemented with 1 mM PMSF and PIC) and sonicated (Bandelin Sonopuls HD2070); Next, proteins were separated by SDS–PAGE, transferred into a nitrocellulose membrane, and stained with primary antibodies (1:5000) at 4°C overnight. After subsequent staining with HRP-conjugated secondary antibodies (1:5000 for antirabbit and 1:2500 for anti-mouse antibodies; room temperature; 2 h), the signal was developed with the SuperSignal™ West Pico Chemiluminescent Substrate and pictures were acquired using ChemiDoc-IT2 (UVP, Meranco, Poznan, Poland). H3 was used as the control.

### 2.11 Confocal Microscopy

For the confocal imaging of proteins, cells were seeded on a Nunc™ Lab-Tek™ chamber slide. 2 h before cells fixation Lysotracker™ Deep Red was added to the final volume of 75 nM, and cells were incubated 1 h at 37°C. Cells were fixed with a 1% formaldehyde solution in PBS at room temperature for 15 min, permeabilized and blocked with 1% FBS solution in PBS with 0.1% TritonX-100 at room temperature for 1h. Primary antibodies (1:400) were added in 1% BSA solution in PBS with 0.1% TritonX-100 and incubated at 4 °C overnight. Next, a secondary antibody (1:400) was added in 1% BSA solution in PBS with 0.1% TritonX-100 at room temperature for 2h. After washing, the slides were mounted with SlowFade™ glass soft-set antifade mountant (with DAPI). TCS SP8 (Leica Microsystems, Germany) with a 63x/1.40 objective (HC PL APO CS2, Leica Microsystems, Germany) was used for sample visualization. The samples were imaged with the following wavelength values for excitation and emission: 485 and 500-550 nm for Alexa Fluor® 488, 550 and 570-580 nm for Alexa Fluor® 546, 480 and 570-580 nm for R-phycoerythrin (PE), 660-670 for Lysotracker Deep Red and 405 and 430-480 nm for DAPI. The fluorescence intensity and colocalization was determined in arbitrary units (a.u.) with Leica Application Suite X (LAS X, Leica Microsystems, Germany). The scans of cells were deconvolved using 3D-Deconwolution accessible in Leica Application Suite X software (LAS X, Leica Microsystems, Germany).

For the visualization of drug accumulation:

1. cells seeded on Nunc™Lab-Tek™chamber slides were treated with anticancer drugs and inhibitors. Next, Lysotracker™ Deep Red (75 nM for 2h), was added to the culture media. After incubation and cell washing with PBS, cells were mounted with SlowFade™ glass soft-set antifade mountant (with DAPI);
2. 3-week old 3D cell cultures were stained with Lysotracker™ Deep Red (75 nM) for 2 h. Next, spheroids was fixed with a 4% formaldehyde solution in PBS, at room temperature for 30 min. After fixation the spheroids were washed with PBS and incubated with 1 µg/ml DAPI for 30 min, at room temperature. Spheroids were analysed immediately.

TCS SP8 (Leica Microsystems, Germany) with a 63x/1.40 and 10x 0.40 DRY objectives (HC PL APO CS2, Leica Microsystems, Germany) was used for sample visualization. The samples were imaged with the following wavelength values for excitation and emission: 485 and 500-550 nm for Alexa Fluor® 488 conjugated Paclitaxel, 470 and 580-600 for Doxorubicin and 405 and 430-480 nm for DAPI. The fluorescence intensity was determined in arbitrary units (a.u.) with Leica Application Suite X (LAS X, Leica Microsystems, Germany).

### 2.12 Resazurin Toxicity Assay

The day prior to transfection/treatment, cells were seeded at a density of 2,500 cells per well on Nunc® MicroWell™ 384-well optical bottom plates or 5,000 cells per well on 96-well plates. After transfection/incubation with drugs and inhibitors, cells were incubated with resazurin solution (5 µM) in the growth medium at 37 °C for 3h. The fluorescence that corresponds to the metabolic activity of living cells was measured with a fluorescence microplate reader (BioTek Synergy HTX, Biokom, Poland) at excitation 530 and emission 590 nm. The fluorescence value for control cells was assumed to be 100%.

### 2.13 Annexin V and propidium iodide staining

For the visualization of apoptosis/necrosis induction by the combination of drugs and inhibitors, Nunc™Lab-Tek™chamber slides were coated with faCellitate BIOFLOAT FLEX coating solution according to the manufacturer protocol. Cells seeded on coated chamber slides were harvested for a 3-weeks to obtain spheroids formation. Next, cells were treated with anticancer drugs and inhibitor. After washing with PBS, according to the manufacturer protocol Annexin V-FITC and Propidium iodide were added to the cells in binding buffer and incubated for 30 min at room temperature. After incubation, spheroids were washed with PBS and incubated with 1µg/ml DAPI for 30 min at room temperature. Spheroids were stored in PBS. Subsequently, for the imaging of the samples, the confocal laser scanning microscopy platform TCS SP8 (Leica Microsystems, Germany) with a 10x objective (HC PL APO CS2, Leica Microsystems, Germany) was used. The samples were imaged with the following wavelength values for excitation and emission 485 and 500-550 nm for AnnexinV-FITC, 450 and 610-620 for Propidium iodide and 405 and 430-480 nm for DAPI, using Leica Application Suite X (LAS X, Leica Microsystems). The level of baseline fluorescence was established individually for each experiment. Fluorescence intensity was determined as the arbitrary units (a.u.) with Leica Application Suite X (LAS X, Leica Microsystems, Germany).

### 2.14 Clinical data analysis

The correlations between SMARCA4 and some ABC gene expression in TCGA Pan-Cancer (PANCAN) dataset of 12 839 samples was explored in UCSC Xena Functional Genomics Explorer. Samples with nulls were removed and data for SMARCA4 expression, which comprised log2(norm_value+1) between 9.6 and 14, were sub-grupped into SMARCA4 low (log2(norm_value+1) < 10.7) and SMARCA4 medium and high (log2(norm_value+1) > 10.7). The threshold was set to include most of deleterious somatic mutation (SNP and INDEL; MC3 public version) in SMARCA4 low subgroup.

### 2.15 Statistical Analysis

Data are shown as mean ± standard deviation (SD). Parametric or non-parametric test was conducted after testing Gaussian distribution of data with the Shapiro–Wilk test. Student’s t test or the Mann–Whitney test was used to calculate statistically significant differences between two samples, while one-way analysis of variance (ANOVA) or Kruskal-Wallis test followed by corresponding post hoc test was carried out to compare multiple samples. Statistics were calculated using GraphPad Prism 8.01 software. Statistically significant differences were marked with * when p < 0.05, ** when p < 0.01, *** when p < 0.001.

## 3. Results

### 3.1 BRG1 is enriched at the promoters of genes functionally linked to endo-lysosomal system and some ABC transporters

Bearing in mind the documented involvement of BRG1 in the transcription of some ABC proteins in 3 triple-negative breast cancer cell lines and ADAADiN prevention of single dose, drug-induced ABC overexpression [24], use was made of the TCGA Pan-Cancer (PANCAN) dataset of RNA and was compared to the mRNA level of SMARCA4 with several ABC genes associated with drug resistance (Fig. 1A). We divided data according to *SMARCA4* expression into 2 subgroups, which corresponded to low and moderate/high expression. As expected, the low mRNA level of SMARCA4 was associated with deleterious mutations in some cancer samples. Additionally, it also correlated with low mRNA levels of *ABCC1*, *ABCC5* and *ABCC10*, but inversely with *ABCB1*, *ABCC2*, *ABCC3* and *ABCG2*. This suggests that BRG1 contributes to the transcriptional regulation of some ABC genes in various cancers.

**Figure 1.**
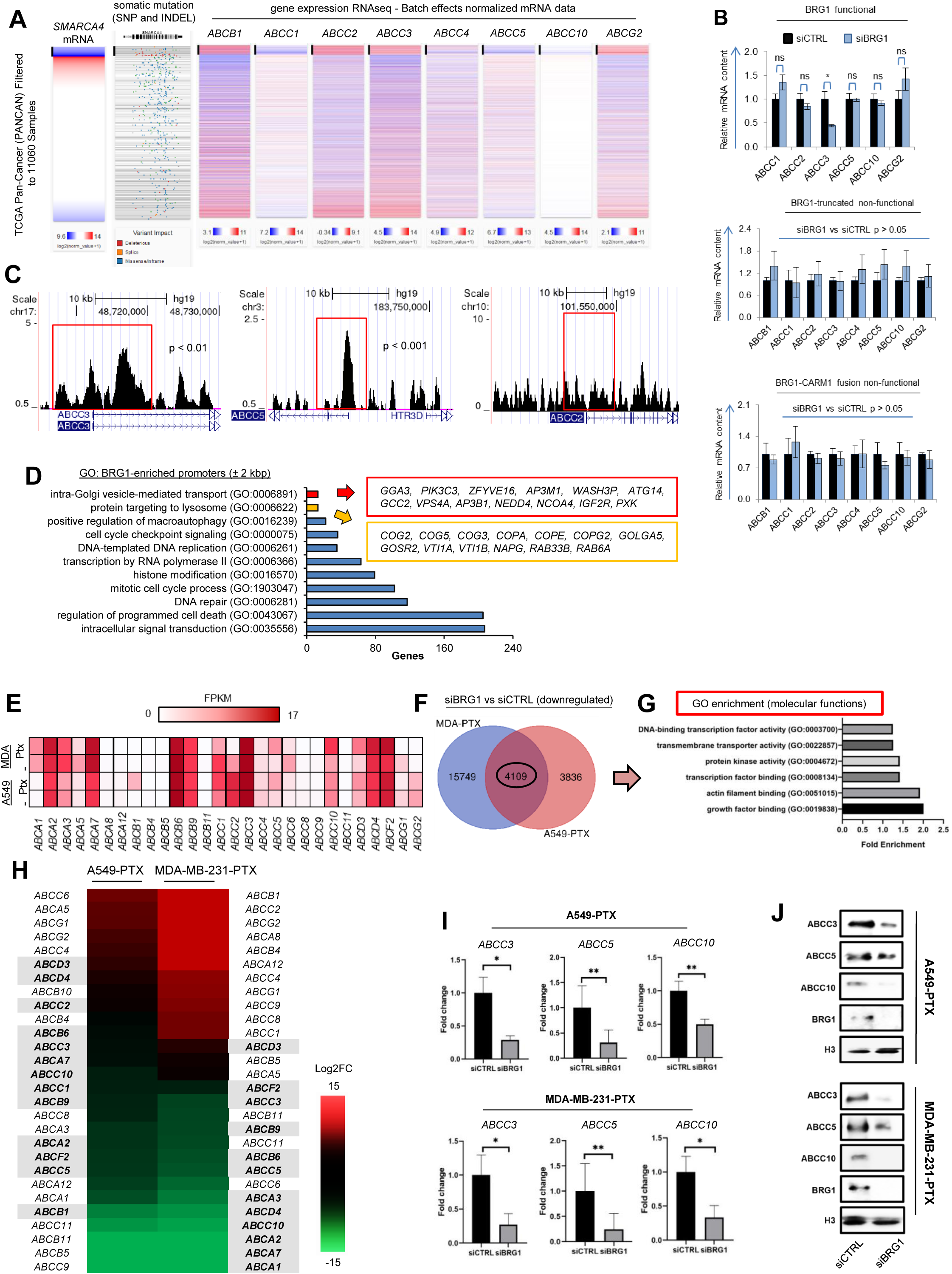
BRG1 co-activates transcription of some multidrug resistance-relevant ABC transporters in paclitaxel-resistant cell lines. (A) mRNA level of *SMARCA4* and its truncating deletions correlate with transcription yield of some transmembrane drug carriers in cancer cells according to clinical data. Visual Spreadsheet of UCSC Xena Functional Genomics Explorer compares co-expression of *SMARCA4* and some multidrug resistance-relevant ABC genes based on TCGA Pan-Cancer (PANCAN). Samples with log2(norm_value+1) < 10.7 were assigned as *SMARCA4* low. (B) Deficiency of BRG1 declines transcription of some ABC genes only in *SMARCA4* wild-type cell lines. The impact of siBRG1 72 h after cell transfection on the studied ABC gene transcription in various *SMARCA4* genotypes was analysed by real-time PCR. Normalized mRNA level of each gene was assumed as 1 in siCTRL. MDA-MB-231 served as *SMARCA4* proficient (BRG1 functional), A549 as *SMARCA4* deficient (BRG1 truncated, non-functional) and MCF7 as *SMARCA4* fusion (BRG1-CARM1 fusion, non-functional). The difference between siCTRL and siBRG1 was analyzed by Student’s t-test with Welsch correction, and “*” when p<0.05, whereas “ns” when p>0.05. (C) BRG1 is enriched at the promoters of ABCC genes, regardless of their transcriptional dependence on BRG1. BRG1 occurrence at the genomic locations spanning TSS of *ABCC3*, *ABCC5* and *ABCC2* in MDA-MB-231 cells was visualized in USCS Genome Browser and BigWig file was derived from MACS2 peak calling. P indicates cutoff for peak detection, when BRG1 was statistically overrepresented at the considered gene promoters. *ABCC2* served as a negative control. (D) BRG1-enriched promoters (± 2 kbp from TSS) represent genes functionally linked to various intracellular processes including protein trafficking to endomembrane system. Ontology of genes characterized by BRG1 occurrence at their promoters (minimum FDR (q-value) cutoff for peak detection set at 0.05) in MDA-MB-231 cells was annotated to biological processes in Panther using statistical overrepresentation test. Genes listed in boxes represent two GO terms (GO:0006891 and GO: 0006622) associated with intracellular vesicle-mediated exchange system of membrane components comprising Golgi apparatus and endolysosomal system. (E) Acquired resistance to paclitaxel changes transcription profile of ABC genes that are functionally linked to multidrug resistance. Normalized gene expression between mapped reads of RNA-Seq datasets from basal (non-resistant; marked as “-“) and paclitaxel-resistant (“Ptx”) cells was generated in CuffLinks using GTEx Gene as a template. Results are shown as a heatmap of normalized gene transcription (FPKM). (F-G) Transient silencing of BRG1 downregulates transcription of a common gene subset, which represent molecular function of transmembrane transporter activity (GO:0022857) in paclitaxel-resistant MDA-MB-251 (MDA-PTX) and A549 (A549-PTX) cell lines. (F) Venn diagram of genes characterized by transcription decline in response to BRG1 silencing (Log2FC < -0.5 ; differential gene expression was generated in CuffDiff using RNA-Seq datasets from BRG1 proficient – siCTRL and deficient – siBRG1 paclitaxel-resistant cell lines) were created using https://bioinformatics.psb.ugent.be/webtools/Venn/ (G) GO terms (molecular function) were assigned to the common gene sets of MDA-PTX and A549-PTX in Panther using statistical overrepresentation test. (H) BRG1 silencing alters transcription of ABC genes, which can contribute to multidrug resistance in cells exposed to several cycles of paclitaxel treatment. Heatmap of differential gene expression presents Log2FC generated in CuffDiff based on RNA-Seq data from BRG1 proficient – siCTRL and deficient – siBRG1 paclitaxel-resistant cells. Bolded genes are characterized by relatively high, unchanged expression or substantial transcription increase caused by paclitaxel. (I) mRNA level of ABC transporters was compared between control and BRG1-deficient samples by real-time PCR. Transcription level was normalized first to housekeeping genes (ACTB, GAPDH and HPRT1) and, then, mRNA level of control was assumed as 1. The difference between two means was tested with Student’s t-test, and statistically significant differences are marked with * when p < 0.05 * and ** when p < 0.01. (J) Effect of transient BRG1 silencing on ABCC3, ABCC5 and ABCC10 protein levels. The lysates of paclitaxel-resistant cells transiently silenced siCTRL and siBRG1 were analyzed by Western Blot. BRG1 was used as a control for silencing efficacy. Histone H3 was used as a loading control.

To confirm the role of BRG1 in the transcription control of the ABC gene subset, the impact of BRG1 transient silencing on the mRNA level in 3 cell lines was tested, this differed in the status of *SMARCA4* (Fig.1B). According to the TCGA database of cancer cell lines, MDA-MB-231 cells carry the wild-type and hence, the fully functional *SMARCA4* transcript. In contrast, the A549 cell line harbors homozygous truncating nonsense mutations leading to BRG1 protein loss [32]. The breast cancer MCF7 line is characterized by the fusion of the *SMARCA4* gene with *CARM1*, making this transcript non-functional [33]. siBRG1 significantly decreased the expression of ABCC3 in cells with the wild-type SMARCA4 but did not cause any substantial difference in cells with SMARCA4 mutations.

In search for the molecular and functional link between BRG1 and the ABC gene transcription in MDA-MB-231, BRG1 ChIP-Seq was performed (Fig. 1C). The MACS2 peak calling revealed BRG1 enrichment at the promoter of *ABCC3* and *ABCC5*, but not *ABCC2*, which remained unchanged in the cells transiently transfected with siBRG1. The occurrence of BRG1 at the promoter of ABCC5 did not render gene transcription dependency on BRG1-SWI/SNF activity as shown in Fig. 1B. BRG1 was found at the promoters of genes annotated to key cellular processes such as replication, transcription, mitotic division, DNA repair, signal transduction, regulation of apoptosis, histone modifications and many others (Fig. 1D) but also for intra-Golgi vesicle mediated transport and protein targeting to lysosomes, which could impact ABC distribution and partitioning within the endolysosomal system and plasma membrane.

### 3.2 BRG1 defines the profile of ABC gene expression in paclitaxel-resistant A549 and MDA-MB-231 cell lines

To identify the ABC genes that can be *bona fide* players in the multidrug resistance of paclitaxel-resistant phenotypes, gene transcription was measured in non-resistant and drug-resistant phenotypes by RNA-Seq, initially assuming A549-PTX as BRG1 deficient and as a control for BRG1-independent gene transcription. For this purpose, normalized (FPKM) counts were juxtaposed for particular mRNAs. To have considerable gene numbers and BRG1-enriched sequences, the list of ABC genes taken into account was extended, all of which have been linked to drug resistance in the literature. As shown in Fig. 1E, both cell lines were characterized by relatively high abundance of *ABCA2*, *ABCA7*, *ABCB6*, *ABCB9*, *ABCC3* mRNA, which responded to cell treatment with paclitaxel with their levels remaining unchanged even when comparing with non-versus paclitaxel-resistant cell lines.

In search for the possible BRG1 contribution to the transcription of ABC genes in paclitaxel-resistant cells, MDA-MB-231 with the wild-type *SMARCA4* and A549 with *SMARCA4* deleterious mutation were used. The induction of cell resistance to paclitaxel and the characteristics of the acquired phenotype was described in Strachowska *et al* [21] and Gronkowska *et al* [14]. Adaptation of the MDA-MB-231 to paclitaxel did not change the expression of BRG1 (Figure S1A-B). Surprisingly, BRG1 mRNA and protein in paclitaxel-resistant A549 cells was found (Figure S1C-D). Sanger sequencing of the *SMARCA4* fragment that spans the c.2184_2206del23 deletion, causing a change in the reading frame, the premature appearance of the STOP codon and lack of protein, indicated the additional deletion of 7 nucleotides and the insertion of 2 nucleotides in the *SMARCA4* sequence before the deletion site (Fig. S1F). The changes of the nucleotide sequence induced by paclitaxel led to reversal mutation that restored the BRG1 expression in paclitaxel-resistant A549 cells. Since the considered mutation did not occur in any of the BRG1 functional domains, the shortened protein bound to chromatin as confirmed by ChIP-qPCR (Fig. S1E). The enzyme was considerably enriched at the promoters of *ABCB1*, *ABCC3* and *ABCC10* in the drug-resistant phenotype of A549 cell line.

To select genes controlled by BRG1, the transcriptomes of BRG1 proficient (siCTRL) and deficient (siBRG1) paclitaxel-resistant phenotypes by RNA-Seq were compared. Differential gene expression showed that ATP-ase regulated the expression of 7,945 genes in A549-PTX, 19,858 genes in MDA-MB-231-PTX, and shared 4,109 genes of BRG1-dependent genes (Fig. 1F). The annotated ontology of the BRG1-controlled genes, common for both resistant cell lines, also indicated a significant portion of genes that encoded proteins involved in active transmembrane transport (Fig. 1G). This group included several ABC family transporters with known roles in resistance to chemotherapeutics such as doxorubicin, paclitaxel, cisplatin, etoposide and methotrexate (Fig. 1H). The number of ABC genes that were highly transcribed in MDA-MB-231-PTX declined in response to BRG1 silencing, whereas the opposite effect was observed for weakly expressed genes. The impact of ATP-ase on ABC expression in A549-PTX was more diverse, but the deficiency of BRG1 was followed by declining transcription of *ABCB1*, *ABCC5*, *ABCC1* and *ABCC10* which are important for drug efflux or sequestration in intracellular organelles. Significantly, BRG1 emerged as transcription co-activator of the 3 genes involved in lysosomal drug trapping,

*ABCC3*, *ABCC5* and *ABCC10*, in both paclitaxel-resistant cell lines. This was further confirmed by real-time PCR and western blot (Fig. 1I-J). In addition, the transcription of *ABCB1* that contributed to lysosomal drug sequestration and increased in response to paclitaxel was also controlled by BRG1 in A549-PTX. Interestingly, silencing of BRM does not reduce the expression of ABC5 and ABCC10 transporters (Fig. S1G). This is all evidence that BRG1 may be crucial for the expression of the ABC genes which protect cells against some anticancer drugs.

### 3.3 BRG1 redistribution in the genome and *de novo* recruitment to gene promoters in the paclitaxel-resistant cell links this enzyme with endolysosomal organization and transport

Cellular adaptation to paclitaxel was followed by BRG1 redistribution in the genome (Fig. 2A-C). Analysis of ChIP-Seq data from non-resistant and resistant MDA-MB-231 cells indicated the recruitment of BRG1 to chromatin since the number of MACS2 computed peaks increased from 15202 to 47231 respectively, with a minimum FDR cutoff for peak detection < 0.001 (Sup. Table 2). However, the proportion of BRG1 enriched gene promoters declined from ∼21% to ∼11%, whereas its binding to enhancers listed in the Vista database increased to ∼0.7%. The observed changes in gene regulatory regions were associated with the shift of 7% ATPase peaks from intergenic to intragenic genome fragments (Fig. 2A and 2B). Gene promoters of paclitaxel-resistant MDA-MB-231 cells were characterized by the altered profile of BRG1 distribution (Fig. 2C and 2D). The considerable extrusion of the enzyme from TSS was followed by peak enrichment in regions between 5 – 20 kbp from the nearest TSS. It may suggest BRG1 enhanced interaction with distal regulatory elements, whereas the depression in BRG1 occurrence at TSS may be linked to the formation of a transcription initiation complex as a consequence of paclitaxel-triggered transcription activation of numerous genes.

**Figure 2.**
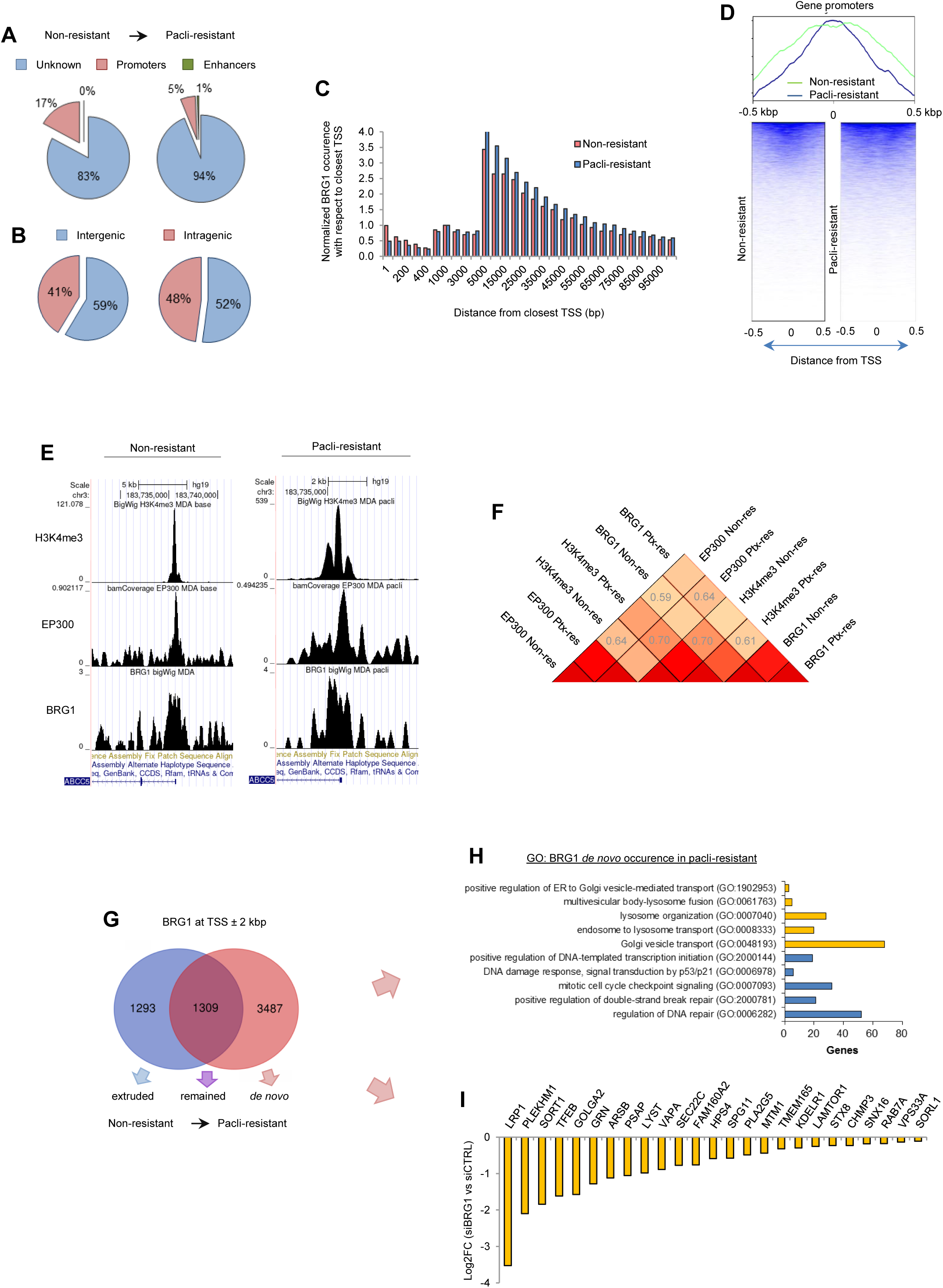
BRG1 redistribution in the genome and de novo recruitment to gene promoters in the paclitaxel-resistant cell links this enzyme with endolysosomal organization and transport. (A-B) Resistance to paclitaxel changes proportion of BRG1 occurrence at the gene promoters. BRG1 peaks were quantified in the following genomic regions: (A) promoters – regions ± 2 kbp from transcription start site (TSS) and enhancers – derived from UCSC table vistaEnhancers, (B) genes (intergenic regions) – derived from UCSC table gtexGene V8. (C) BRG1 shifts from TSS proximal to distal regions in response to repeated MDA-MB-231 cell exposure to paclitaxel. BRG1 peaks were quantified with respect to the TSS by bedtools ClosestBed, while taking MACS2 peak summits of BRG1 and TSS derived from UCSC table gtexGene V8. Counts were normalized to peak number at – 1 kbp. (D) Gained resistance to paclitaxel is associated with BRG1 spreading at the gene transcription start site. BRG1 occurrence in the region spanning TSS (± 0.5 kbp) was monitored by plotting a profile and heatmap of BRG1 peaks against TSS as a reference point (computeMatrix: plotHeatmap; regions sorted in descending order by mean without clustering). (E) BRG1, EP300 and H3K4me3 are enriched at the promoter of *ABCC5* gene regardless of MDA-MB-231 cell resistance to paclitaxel. Genome coverage (as BigWig derived from -bamCoverage) with the two proteins and histone modification around TSS of *ABCC5* was visualized in USCS Genome Browser. (F) BRG1 and EP300 are less centered at the transcriptionally active TSS in paclitaxel-resistant cells. The distribution of these two enzymes at H3K4m3 positive TSS in MDA-MB-231 non-versus paclitaxel-resistant was counted (-computeMatrix) and visualized (-plotHeatmap) in the region spanning TSS ± 2.5 kbp. Regions were clustered with respect to H3K4me3 intensity. (G) resistance to paclitaxel declines BRG1 corelation with H3K4me3, but enhances co-occurrence with EP300. BRG1, EP300 and H3K4me3 genomic co-distribution was calculated as Spearman correlation coefficient (-multiBigWigSummary and - plotCorrelation; genomic regions were taken as H3K4me3 peaks that overlap TSS). (H) Paclitaxel resistance of MDA-MB-231 is associated with substantial recruitment of BRG1 to new subset of gene promoters. Gene promoters (TSS ± 2 kbp) characterized by BRG1 peaks in non-resistant and paclitaxel-resistant phenotypes were plotted in venn diagram. Extruded, *de novo* and remained promoters refer to promoters enriched BRG1 in non-resistant cells only, in paclitaxel-resistant cells, and in both cell phenotypes, respectively. The possible enzyme shift in the considered regions were not taken into consideration. (I) BRG1 is recruited *de novo* to promoters of genes, which are functionally linked to numerous processes including ER-Golgi-endosome-lysome transport. Gene ontology (GO: biological process complete) of BRG1 *de novo* enriched promoters in paclitaxel-resistant MDA-MB-231 cells was assigned in Panther using statistical overrepresentation test

BRG1 co-occurred with EP300 at the promoters of some transcriptionally active genes, as described in previous papers [22,23] (Fig. 2D-4E). These were assigned as genomic fragments spanning TSS ± 2 kbp and were hallmarked by the trimethylation of H3K4. Interestingly, the promoter of ABCC5, which is transcriptionally controlled by BRG1 only in the drug-resistant phenotype, was similarly enriched in BRG1 and EP300 and characterized by a similar profile of H3K4 trimethylation in both cell conditions (Fig. 2D). This suggests that transcription dependence on BRG1 may be determined by another, still unknown factor. As mentioned previously, this can be linked to the reorganization of proximal promoters during gene activation and loading of transcription initiation factors. The Spearman correlation coefficient for BRG1 co-distribution with H3K4me3 declined from 0.7 to 0.61 during the acquisition of drug resistance, suggesting chromatin extrusion or translocation to more distal genomic regions as shown in Fig. 2C. However, the co-occurrence with EP300 increased from 0.59 to 0.64, possibly to facilitate EP300-driven gene transcription (Fig. 2F).

Although the relative BRG1 occurrence declined at the gene promoters that were normalized to total peak number, the number of BRG1-enriched promoters was higher in the paclitaxel-resistant phenotype (Fig. 2G). Annotation of ontology to the genes controlled by promoters (that were characterized by de novo BRG1 enrichment after numerous cell exposures to paclitaxel), disclosed processes that were involved in cell response to genotoxic stress, endosome to lysosome transport, lysosome organization and ER to Golgi vesicle-mediated transport (Fig. 2H-2I). The latter processes can contribute to the observed trafficking of ABCC3, ABCC5 and ABCC10 to lysosomes in paclitaxel-resistant phenotypes. Differential expression of genes transcribed in BRG1 proficient and deficient paclitaxel-resistant MDA-MB-231 cells confirmed the role of ATP-ase in the transcription enhancement of numerous genes assigned to endolysosomal GO terms (Fig. 2I).

### 3.4 BRG1 guides drug distribution to lysosomes and declines paclitaxel-resistant cell sensitivity to drugs

Bearing in mind that BRG1 controls the transcription of ABC genes which were found to be substantially enriched in the lysosomes of paclitaxel-resistant cells, BRG1 deficiency in the decline of ABCC3, ABCC5 and ABCC10 abundance in these organelles was tested by using the immune staining of ABCCs and their confocal imaging against lysosomes and DNA in cells with normal (siCTRL) and knockdown BRG1 (siBRG1). The silencing of BRG1 caused visible and significant reduction of ABCC3 and ABCC10 levels inside cells (Fig. 3A, 3B), and reduced their lysosomal localization, quantified by measuring protein colocalization with lysosomes (Fig. 3C). No changes in the expression or intracellular localization of these proteins were observed after BRM silencing (Fig S2A). It was significant that the BRG1 level had no impact on lysosome content in the paclitaxel-resistant cells (Fig. S2B). Confocal microscopy with a deconvolution algorithm allowed the visualization of the decline in ABCC5 level in the lysosome membrane that was caused by BRG1 silencing in both paclitaxel-resistant cell lines (Fig. 3D).

**Figure 3.**
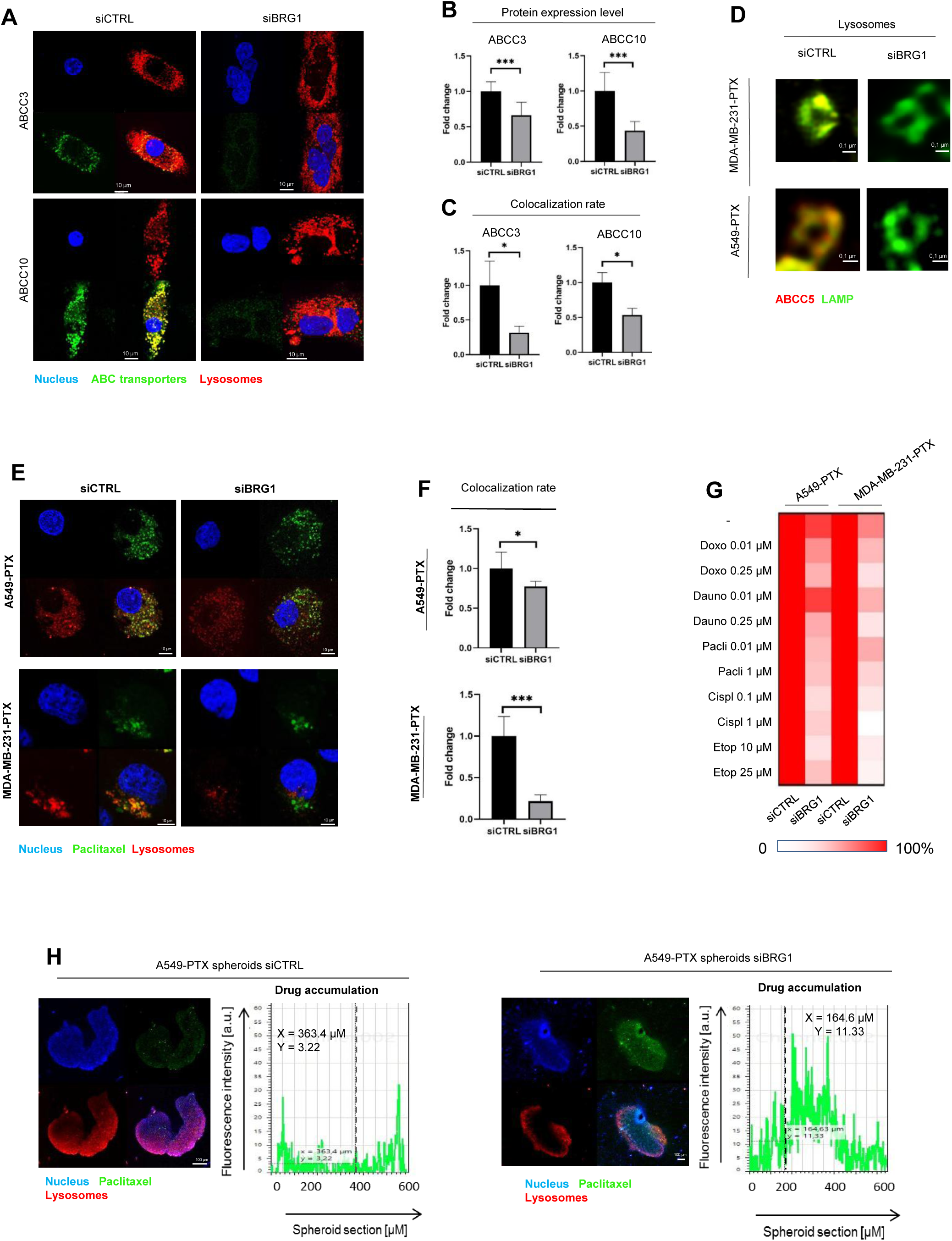
BRG1 drives overexpression of lysosomaly localized ABC-transporters in PTX-resistant cell lines, which are crucial for drug sequestration and reduced drug cytotoxicity. (A-D) BRG1 confers overrepresentation of some ABCC proteins in lysosomes of paclitaxel-resistant cancer cells. (A) ABC transporter expression and localization 72 h after cell transfection with siCTRL and siBRG1 were visualized by immunocytostaining followed by confocal microscopy. Green fluorescence derived from Alexafluor488-conjugated secondary antibody corresponds to ABC transporters appearance in cells. DNA was stained with DAPI (blue). Lysosomes was stained with LysoTracker (red). The fluorescence intensity (B) and colocalization (C) was determined in arbitrary units (a.u.) with Leica Application Suite X. The difference between two means was tested with Student’s t-test, and statistically significant differences are marked with * when p < 0.05 *, ** when p < 0.01, *** when p < 0.001 (D) Red fluorescence of ABC transporters is derived from R-phycoerythrin-labelled secondary antibody and LAMP1-green fluorescence is derived from Alexafluor488-conjugated secondary antibody. The scans of lysosomes were deconvolved using 3D-Deconwolution accessible in Leica Application Suite X software (LAS X, Leica Microsystems, Germany). (E) BRG1 causes lysosomal sequestration of paclitaxel Oregon Green in lysosomes of paclitaxel-resistant cancer cell lines. Colocalization of fluorescently labelled paclitaxel and lysosomes was compared between BRG1 proficient and deficient cells. Paclitaxel is marked in green (Oregon Green 488), lysosomes in red (LysoTracker Deep Red), DNA in blue (DAPI). The colocalization (F) was determined in arbitrary units (a.u.) with Leica Application Suite X. The difference between two means was tested with Student’s t-test, and statistically significant differences are marked with * when p < 0.05 *, ** when p < 0.01, *** when p < 0.001 (G) BRG1 confers multidrug resistance in cells exposed to several doses of paclitaxel. Sensitivity to doxorubicin, daunorubicin, paclitaxel, cisplatin and etoposide was compared between control and BRG1 silenced ptx-resistant cell lines. Viability was measured with resazurin-based assay. Metabolic activity of control cells was assumed as 100%. (H) Paclitaxel Oregon Green accumulation in lysosomes of paclitaxel-resistant cells grown in 3D cultures requires BRG1. Control and BRG1 silenced paclitaxel-resistant A549 cell line spheroids were scanned for intracellular paclitaxel Oregon Green distribution. The drug is marked in green, DNA in blue, lysosomes in red (staining as described in E). The fluorescence intensity plot at spheroid cross section was determined in arbitrary units (a.u.) with Leica Application Suite X.

**Figure 4.**
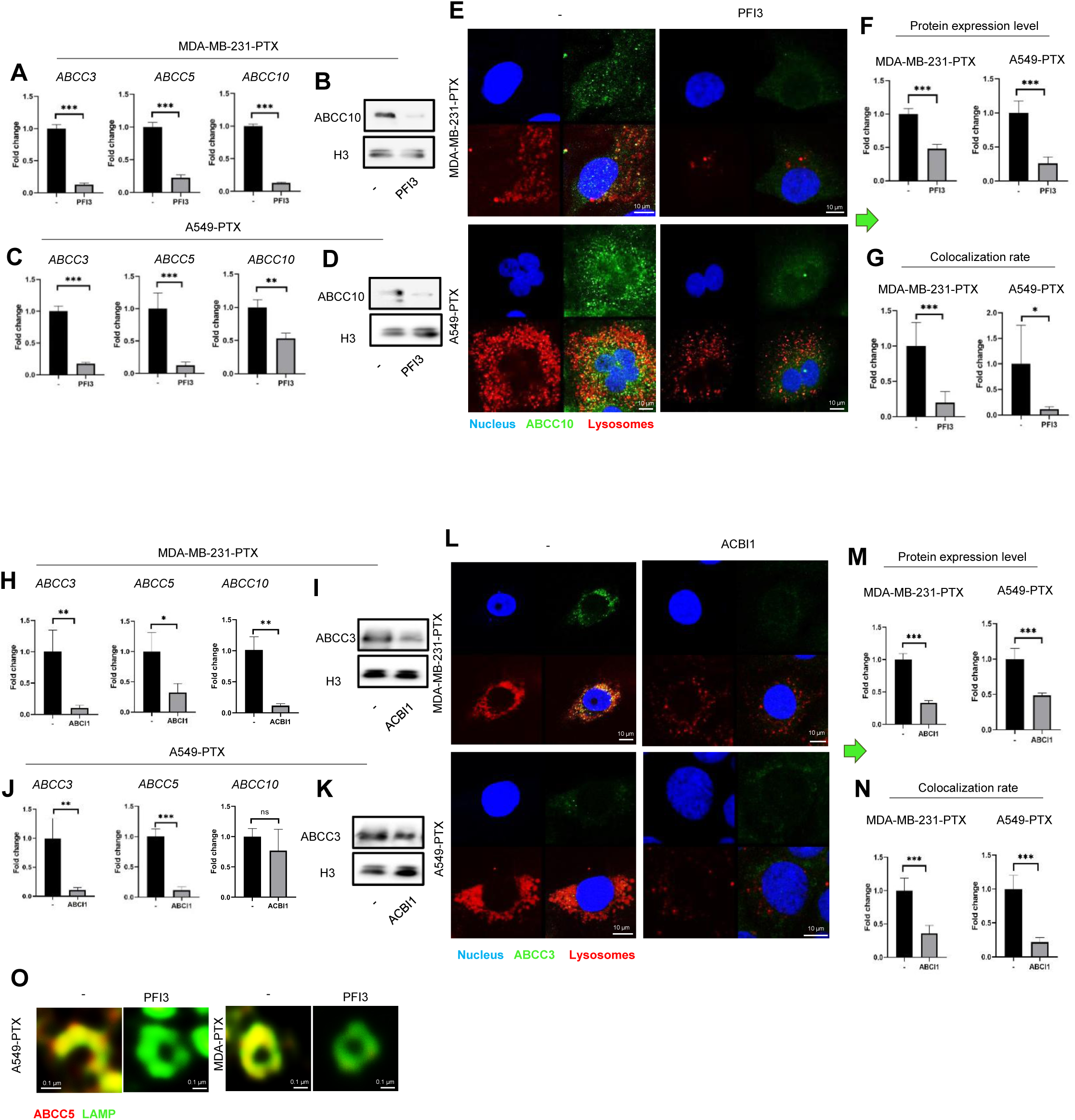
SWI/SNF targeting with PFI-3 and ACBI1 reduces expression on ABCC transporters overrepresented in lysosomes of paclitaxel-resistant cells. (A-D, H-K) The lack of SWI/SNF activity declines transcription and intracellular abundance of ABCC3, ABCC5 and ABCC10 in paclitaxel-resistant phenotypes. mRNA level of *ABCC3*, *ABCC5* and *ABCC10* was compared by real-time PCR in paclitaxel-resistant MDA-MB-231 (A) and A549 (C) cells exposed and not to PFI3 (2.5 uM, 72h) and paclitaxel-resistant MDA-MB-231 (H) and A549 (J) cells exposed and not to ACBI1 (0.5 uM, 72h). Transcription level was normalized first to housekeeping genes (*ACTB*, *GAPDH* and *HPRT1*) and for control sample was assumed as 1. The difference between two means was tested with Student’s t-test, and statistically significant differences are marked with * when p < 0.05 *, ** when p < 0.01, *** when p < 0.001. (B,D) Impact of PFI3 on ABCC10 protein level in paclitaxel-resistant MDA-MB-231 (B) and A549 (D) cell lysates was tested by Western Blot. (I,K) Effect on ACBI1-targeted SWI/SNF subunits degradation on ABCC3 protein level in paclitaxel-resistant MDA-MB-231 (I) and A549 (K) cell lysates was tested by Western Blot. Histone H3 was used as a loading control. (E, L) Expression and localization of ABC transporters was visualized by immunocytostaining followed by confocal microscopy in non-treated vs PFI3 (E) and ACBI1 (L) -targeted cells. Green fluorescence of ABC transporters derived from Alexafluor488-conjugated secondary antibody, blue fluorescence of DNA from DAPI, whereas lysosomal red fluorescence from LysoTracker Deep Red. The fluorescence intensity (F,M) and colocalization (G,N) was determined in arbitrary units (a.u.) with Leica Application Suite X. The difference between two means was tested with Student’s t-test, and statistically significant differences are marked with * when p < 0.05 *, ** when p < 0.01, *** when p < 0.001. (O) Confocal microscopy imaging of lysosomal membrane proteins in control and PFI3-treated MDA-MB-231-PTX cells. ABC transporters were visualized by immunocytostaining followed by confocal microscopy. Red fluorescence of ABC transporters is derived from R-phycoerythrin-labelled secondary antibody and green fluorescence of LAMP1 is derived from Alexafluor488-conjugated secondary antibody. The scans of lysosomes were deconvolved using 3D-Deconwolution accessible in Leica Application Suite X software (LAS X, Leica Microsystems, Germany).

To check the functional impact of BRG1-dependent expression of the ABC transporters which are enriched in lysosomes of paclitaxel-resistant cells on intracellular drug distribution, we compared the colocalization of doxorubicin and paclitaxel OregonGreen with lysosomes in BRG1 proficient and deficient cells using confocal microscopy (Fig. 3E, Fig. S2C). Silencing of the enzyme visibly declined the drug accumulation in lysosomes, quantified as colocalization of drug and lysosome fluorescence (Fig. 3F, Fig. S2D). Previously documented roles of ABCC3, ABCC5 and ABCC10 in lysosome drug accumulation [14] led to the conclusion that the BRG1-dependent transcription of ABCC3, ABCC5 and ABCC10 was responsible for the distribution of their substrates into lysosomes in both paclitaxel resistant breast and lung cancer cells. This also led to the hypothesis that BRG1 may affect the cell response to chemotherapeutics and cause lysosome associated chemoresistance. This was tested by comparing the viability of cells which differed in BRG1 abundance and were exposed to chemotherapeutics of various acidity. This feature of the tested drugs defined the mode of their passive or ABC-dependent influx to lysosomes. The silencing of BRG1 sensitized cells of both lines to all chemotherapeutics tested (Fig. 3G), increased considerably with these acidic drugs that do not enter lysosomes by passive diffusion.

The impact of BRG1 on drug penetration was also tested in cell spheroids. 3D cultures of MDA-PTX and A549-PTX cells were characterized by the accumulation of doxorubicin and paclitaxel OregonGreen in the peripheral layer, whereas BRG1 silencing resulted in drugs reaching the inner spheroid layers (Fig. 3H, S2E). This suggests that BRG1-dependent drug trapping in lysosomes protects the deeper parts of the spheroids from drug cytotoxicity. This conclusion is supported by a previous finding, where lysosome neutralization or inhibition of ABCC reduced spheroid penetration by doxorubicin and paclitaxel OregonGreen as well as increasing the overall number of apoptotic cells inside these spheroids [14].

### 3.5 SWI/SNF targeting with PFI3 and ACBI1 decreases lysosome-associated ABCC in PTX-resistant cancer cell lines

Since BRG1 is responsible for the overexpression of lysosomal pool of ABC transporters and increased chemoresistance of paclitaxel-resistant cells, the BRG1/BRM inhibitor - PFI3 and SMARCA2/4 PROTAC degrader - ACBI1 were tested for their likely potential to sensitize paclitaxel-resistant cell to chemotherapy. The ability of ACBI1 to induce degradation of BRG1 was confirmed by Western Blot (Fig S3A). SWI/SNF inhibitor was found potent to decline both the mRNA and protein level of the three considered multidrug resistance (MDR) transporters (Fig. 4A-D). However, BRG1/BRM degradation with ACBI1 reduced expression of ABCC3 and ABCC5 (Fig. 4H-K), but increased ABCC10. The reduced co-occurrence of ABCC10 and ABCC3 with lysosomal marker (Fig 4 E-G, 4L-N, Fig. S4 A-B Fig. S4 D-E) after PFI3 and ABI1, as well as ABCC5 with LAMP1 (Fig 4O) after PFI3 was visualized and quantified by confocal microscopy in paclitaxel-resistant lines (Fig. 4E-F).

In non-resistant MDA-MB-231 and A549 only highly expressed ABCC3 was declined after cell treatment with PFI3 and ABI1, respectively (Fig. S3 B-E).

PFI3 and ABCI1 altered the intracellular distribution of doxorubicin and paclitaxel OregonGreen by retaining a substantial drug concentration outside lysosomes. This was confirmed by measuring the drug accumulation and drug-lysosome colocalization (Fig. 5A-F, S5A-C). The SWI/SNF targeting molecules also slightly elevated the total drug accumulation inside paclitaxel-resistant cells. Particularly, PFI3 phenocopied the effect of BRG1 silencing on lysosomal delocalization of paclitaxel OregonGreen and doxorubicin (Fig. 3E-F Fig. S2B-C). PFI3 and ABI1 allowed also deeper drug penetration in spheroids, whereas intact 3D cultures were characterized by drug accumulation inside outer cell layers (Fig 5G).

**Figure 5.**
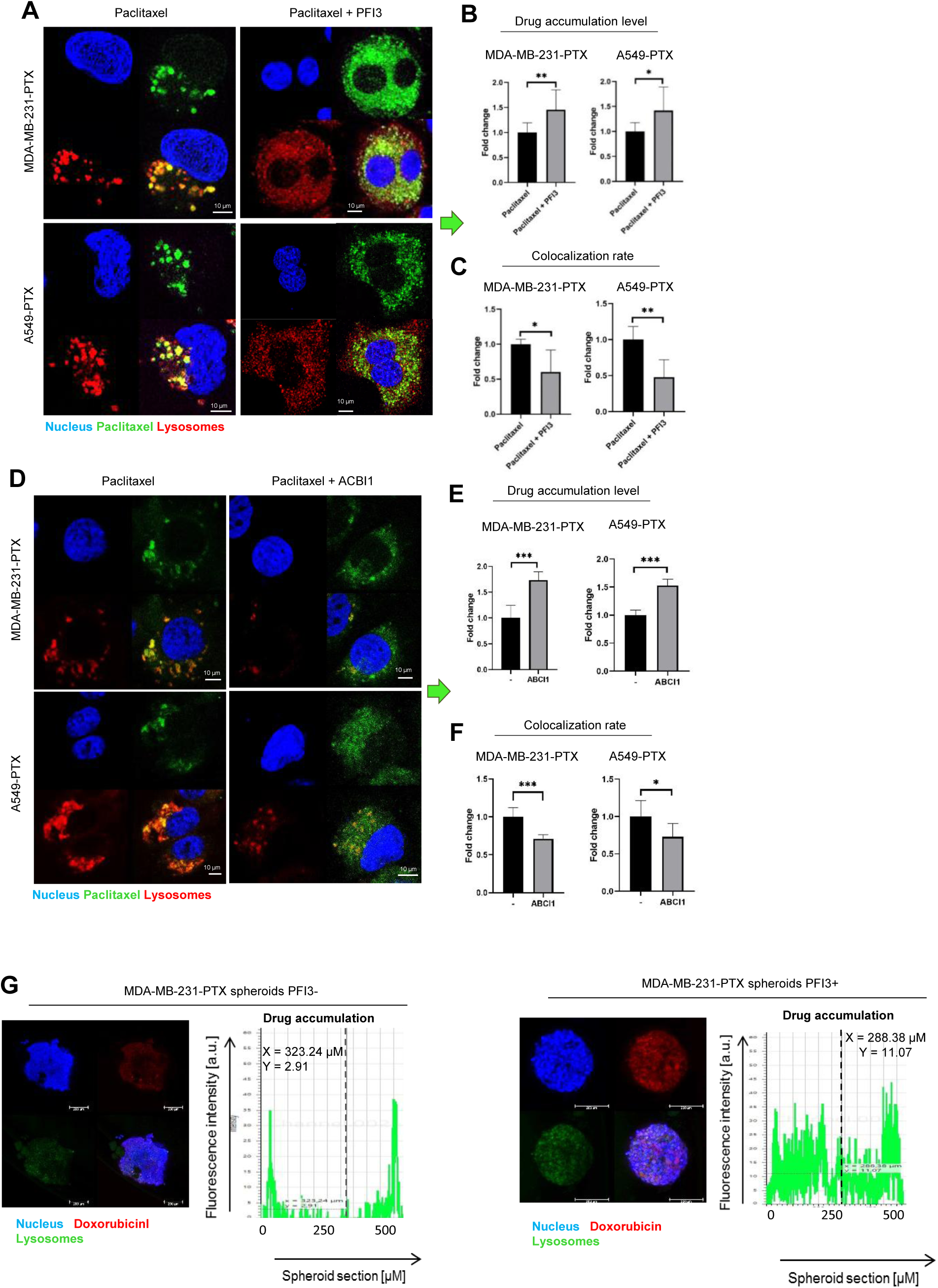
SWI/SNF targeting with PFI-3 and ACBI1decreases drug accumulation in lysosomes of paclitaxel-resistant cancer cells. (A,D) SWI/SNF-targeting uncouples paclitaxel Oregon Green accumulation in lysosomes of paclitaxel-resistant cells. Confocal imaging was used to study the impact of SWI/SNF inhibitor – PFI3 (2.5 uM, 24h) (A) and ABCI1 (D) on intracellular localization of fluorescently labelled paclitaxel. The drug is marked green (Oregon Green 488), DNA in blue (DAPI) and lysosomes in red (LysoTracker Deep Red). The fluorescence intensity of paclitaxel Oregon Green (B, E) and colocalization between the drug and lysosomes (C,F) was determined in arbitrary units (a.u.) with Leica Application Suite X. The difference between two means was tested with Student’s t-test, and statistically significant differences are marked with * when p < 0.05 *, ** when p < 0.01, *** when p < 0.001 (G) Doxorubicin accumulation in lysosomes of paclitaxel-resistant cells grown in 3D cultures after PFI3-treatment. Control and inhibitor-treated paclitaxel-resistant MDA-MB-231 cell line spheroids were scanned for intracellular Doxorubicin distribution. The autofluorescent drug is marked in red, DNA in blue, lysosomes in green. The fluorescence intensity plot at spheroid cross section was determined in arbitrary units (a.u.) with Leica Application Suite X.

### 3.6 PFI3 and ACBI1 augments the toxicity of anticancer drugs in PTX-resistant cancer cell lines

The observed increase in the drug concentration inside cells and the impairment of drug sequestration in lysosomes could be responsible for the altered cytotoxicity of anticancer drugs in SWI/SNF-treated cancer cells in 3D (Fig.6A) and 2D cultures (Fig 6D). In spheroids, the combination of doxorubicin or paclitaxel with PFI3 and ABCI1 was more potent in apoptosis induction than drugs alone as shown in the confocal images and quantification of Annexin V green fluorescence (Fig. 6A-C). Importantly, Annexin V-positive cells were equally distributed across all depths of the spheroids that were exposed to the combination of PFI3 or ABI1 and drugs, whereas apoptotic cells were localized in the peripheral layers similarly to drugs in cells proficient in SWI/SNF activity. The significant induction of cell necrosis was only observed in spheroids exposed to doxorubicin, and further intensification of necrotic cell death as well as apoptotic commitment was caused by ABI1. On the contrary, PFI3 substantially enhanced apoptosis in spheroids as well as in 2D culture treated with paclitaxel (Fig. 6D). PROTAC degrader sensitized 2D model of paclitaxel-resistant cancer cells especially to doxorubicin but also to lower doses of cisplatin and etoposide when added 48 h prior to chemotherapeutics (Fig.6E). SWI/SNF inhibitor moderately increased drug toxicity in the culture of adherent cells (Fig.6F).

**Figure 6.**
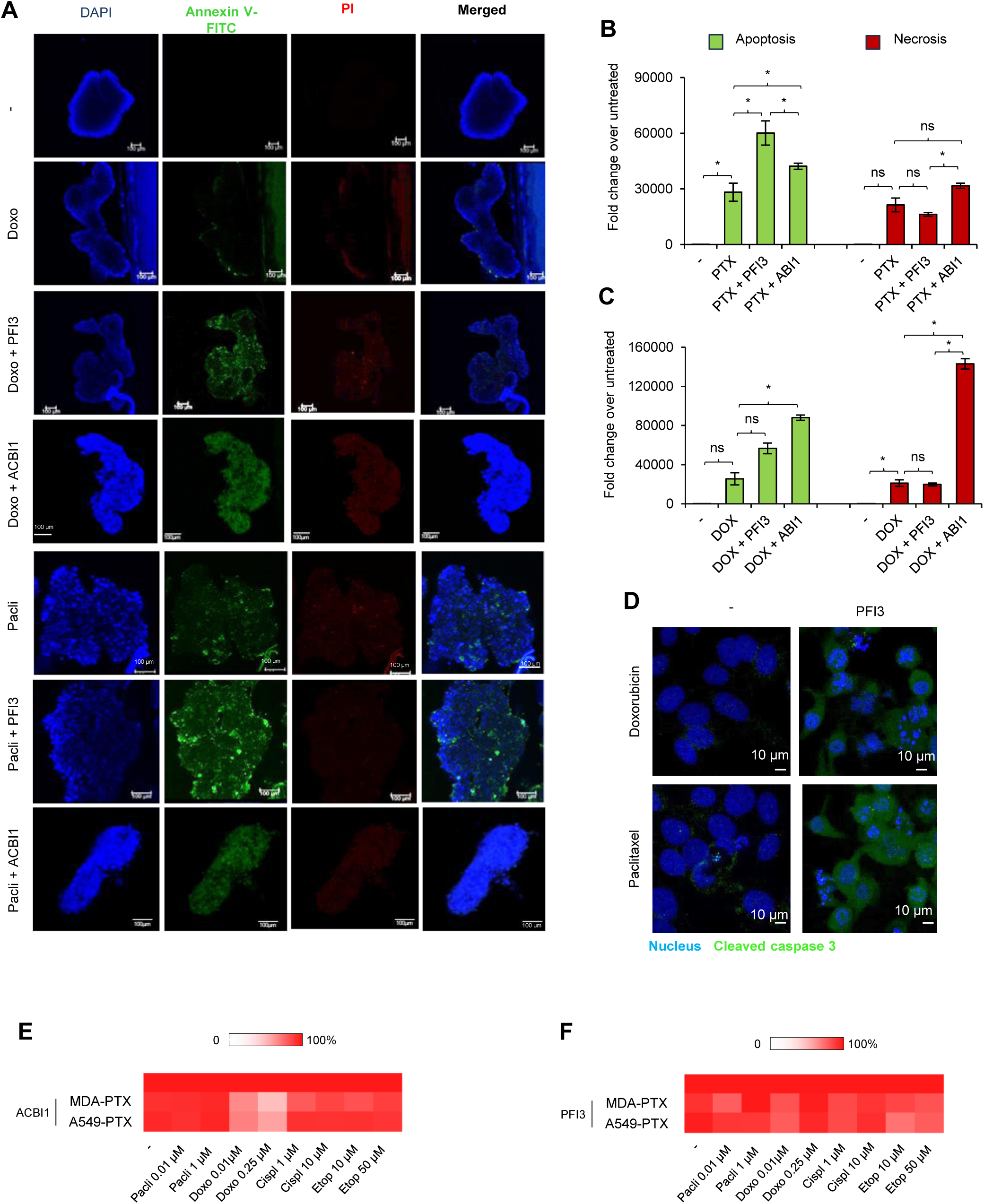
PFI3 and ACBI1 augments the toxicity of anticancer drugs in PTX-resistant cancer cell lines. (A) SWI/SNF-targeting increases paclitaxel-resistant cell death in a spheroid model of anticancer treatment. 3-week spheroids were firstly incubated with PFI3 or ABCI1 for 72 h and then paclitaxel and doxorubicin were added for 48 h. Externalization of phosphatidylserine, which marks apoptotic cells, was monitored by confocal microscopy after cell staining with FITC-conjugated annexin V (green). DNA of necrotic cells (with disrupted membrane integrity) was stained with Propidium Iodide (red). DNA was stained with DAPI (blue). (B, C) The fluorescence intensity was determined in arbitrary units (a.u.) with Leica Application Suite X. The difference between two means was tested with Student’s t-test, and statistically significant differences are marked with * when p < 0.05 *, ** when p < 0.01, *** when p < 0.001. (D) PFI3 (2.5 uM, 72h) enhances apoptosis induced doxorubicin and paclitaxel in A549-PTX cells. Confocal imaging using Leica TCS SP8 (Leica Microsystems, Germany) was used to compare the intensity of caspase 3 fluorescence between PFI3-treated and non-treated cells. The impact of ACBI1 (0.5 µM, 48 h) (E) and PFI3 (2.5 µM, 48 h) on the toxicity of some anticancer therapeutics. Cell viability was quantified with resazurin-based assay 24 after cell treatment with drugs. Cell viability with drug alone was assumed as 100%.

All these results indicate that SWI/SNF targeting is effective for sensitizing paclitaxel-resistant cancer cells to chemotherapeutic agents for the 2 studied cells lines. This may be attributed to the reduced expression of ABC transporters and prevention of the lysosomal trapping of the drugs. From clinical perspective the combination of ABI1 with doxorubicin seems worth of further investigation for the treatment of paclitaxel-resistant triple-negative breast cancers.

## 4. Discussion

The expression profile of resistance proteins, which are responsible for the failure of chemotherapy in cancer treatment, can change with disease progression [34]. The epithelial-mesenchymal transition associated with tumor progression has been shown to increase the expression of ABC transporters [35]. Due to the important role of ABC transporters in the occurrence of this phenomenon, they are a target for the development of combined therapies aimed at sensitizing cancer cells to standard chemotherapeutics. Overcoming multidrug resistance, which is associated with the expression of ABC transporter family proteins, is a challenge and none of the 3 generations of ABC inhibitors, (particularly in the case of ABCB1), have provided satisfying results in clinical trials and all have failed to become FDA approved drugs. Our current study indicates that inhibition of SWI/SNF with PFI3 or degradation of SWI/SNF ATPases with PROTAC allows to substantially increase drug toxicity due to simultaneous reduction of several ABC transporters in paclitaxel-resistant phenotypes of the chosen lung and breast cancers. SWI/SNF targeting increased overall drug accumulation may be due to reduced drug uptake by lysosomes associated with a decrease in lysosomal ABC transporters and thus reduced exocytosis of chemotherapeutic drug-loaded lysosomes [36]. Another explanation is the inhibition of the expression or activity of drug transporters located in the plasma membrane of paclitaxel-resistant phenotypes, such as *ABCC1* in A549-PTX. Since these compounds target both ATPases, some genes related to drug transport may be upregulated by BRM and thus downregulated in response to SWI/SNF targeting. These finding are particularly important in light of studies, which document the scarce sensitizing impact of single inhibition of ABCB1, ABCG2 or some of the ABCC subfamily to chemotherapeutics and poor drug accumulation inside drug-resistant cells [21].

The identification of an upstream mechanism responsible for transcription of several *ABC* genes under chemotoxic stress could provide new targets to limit the drug efflux from resistant cancer cells and, hence, a new approach for combined and improved chemotherapy. Experimental evidence is provided here that considers the role of BRG1 in transcriptional control of ABC genes that are overexpressed in paclitaxel-resistant cell lines. Specifically, the focus was on the impact of BRG1 on *ABCC3*, *ABCC5* and *ABCC10*, which are enriched to lysosomes in paclitaxel-resistant cells, and the last 2 genes that are overexpressed once the cell is exposed to several doses of paclitaxel. The functional analysis of these 3 proteins that occur in lysosomes, disclosed their role in the sequestration of doxorubicin and paclitaxel OregonGreen, hence these membrane transporters are capable of limiting drug cytotoxicity. The active promoters of these genes are characterized by H3K4 trimethylation and occurrence of BRG1 and EP300. Such a complex previously described by us and others is capable of chromatin decondensation and defines transcription permissive environment because of extrusion of acetylated nucleosomes [22]. The functional interaction between BRG1 and EP300 can be also confirmed by the fact that EP300 bromodomain inhibitor I-CBP112 substantially decline transcription of ABC genes in drug-resistant cell lines. Furthermore, development of resistance to paclitaxel increased the number of BRG1-bound promoters of genes linked to endolysosomal and ER-Golgi protein transport, which may also define the profile of ABC proteins in lysosomes and their enrichment in these organelles. Although BRG1 was *de novo* recruited to the promoter of *TFEB* gene, which encodes the master regulator of lysosome biogenesis during MDA-MB-231 cell adaptation to paclitaxel and the BRG1 silencing in resistant cells that caused substantial decline in TFEB gene transcription, the number of lysosomes estimated in confocal images remained unchanged [37]. This can be explained by the fact that regulation of TFEB occurs predominantly by post-translational modifications such as phosphorylation, acetylation, SUMOylating, PARylation, and glycosylation, which allowed the cell to respond quickly to nutrient fluctuations but also to stress [38]. And the adaptation-induced increase in TFEB abundance may not determine further enhance in lysosome biogenesis. Moreover, other BRG1-dependent genes that regulate endolysosomal pathways need to be taken into consideration while explaining this drug trapping in lysosomes. In particular, the mechanism that drives ABCC5 and ABCC10 trafficking to lysosomes and ABCC3 redistribution to these organelles in paclitaxel-resistant cells remains unknown. This was discussed in a previous paper [14], where the topological inversion of the plasma membrane ABCB1 via endocytosis resulting in the transporter actively pumping agents into lysosomes was proposed. This may also be true for ABCC3, but neither ABCC5 nor ABCC10 was detected in the plasma membrane of the non-resistant or paclitaxel-resistant cells. Hence, the protein sorting from Golgi to the endolysosomal system could be taken into consideration. Although N-acetylglucosamine-1-phosphodiester α-N-acetyl-glucosaminidase declined in response to BRG1 silencing, the other 2 crucial elements of the mannose-6-phosphate (M6P) pathway that is responsible for the transport of hydrolytic enzymes to lysosomes: the Mannose-6 phosphate receptor and GlcNAc-1-phosphotransferase (GNPTAB and GNPTG), were differently regulated by BRG1 in paclitaxel-resistant cell lines. Furthermore, all 3 GGAs (Golgi-localized, γ-ear-containing, Arf (ADP-ribosylation factor)-binding proteins), which are a family of ubiquitously expressed, Arf-dependent, clathrin adaptors responsible for the sorting of mannose-6-phosphate receptors (MPRs) between the trans-Golgi network (TGN) and endosomes, declined considerably in BRG1-deficient cells. The mechanism of lysosome enrichment in ABCC3, ABCC5 and ABCC10 remains unsolved and needs further experimental verification.

In line with our study, some attempts have already been made to use BRG1-targeted therapies to sensitize cancer cells to chemotherapy. PFI-3 exerted a DNA damage– sensitizing effect by directly blocking SWI/SNF’s chromatin binding, leading to defects in DSB repair and aberrations in the damage checkpoints in A549 and HT29 cells. This resulted in the increase of cell death primarily via necrosis and senescence after doxorubicin treatment [39]. Furthermore, PFI-3 and Structurally Related Analogs of PFI-3 (SRAPs) sensitized LN229 glioblastoma cells and patient-derived glioblastoma cells: GBM6, GBMX10, and GBMX16, to chemotherapeutic drugs such as temozolomide and carmustine *via* changes in the expression of genes implicated in the glucose metabolism [40,41]. It was also showed that the reduction of BRG1 activity by using ADAADi (Active DNA-dependent ATPase A Domain inhibitor) increased the chemosensitivity of MDA-MB-231 cells to 5-fluorouracil, cisplatin, cyclophosphamide, doxorubicin, gemtabicine and paclitaxel by reducing the expression of ABC transporters and preventing their overexpression induced by a single dose of anticancer drugs [24]. Not much is known about the pharmacokinetics of PFI-3 and ADAADi *in vivo*, so their possible application in anticancer therapies will require further investigation. Importantly, these 2 compounds are not selective for BRG1, hence their use should be considered in a context-specific manner. PFI-3 targets the bromodomains of BRG1 and BRM, whereas ADAADi interacts with ATP-dependent chromatin remodeling proteins through a motif that is present in the conserved helicase domain. However variable responses were observed in different cell lines [42]. BRG1-specific effects of PFI-3 on lysosomal drug sequestration and ABCC enrichment in these organelles in paclitaxel-resistant cells was confirmed by BRG1 silencing, as well as by very low or no-impact of these compounds on the studied parameters in non-resistant cells. Hence, PFI-3 and other possible BRG1/BRM inhibitors emerged as candidates for anticancer approaches combined with chemotherapy drugs in the subset of BRG1-proficient cancers that developed multidrug resistance by BRG1-dependent drug trapping in lysosomes. Even less is known about pharmacokinetics and possible side effects of SMARCA2, SMARCA4 and PBRM1 degrader – ACBI1, which emerged here as potent interrupter of multidrug resistance in the studied cell lines. Former papers described anti-proliferative and pro-apoptotic activity of this compound in the culture of SK-MEL-5 melanoma cells [43]. In light of our findings this compound can be considered for an anticancer treatment modality that combines two or more therapeutic agents. Lysosome targeted drug combinations have been tested recently *in vitro* due to the role that these organelles have shown in passive and active trapping of chemotherapeutics [44]. Numerous chemical agents and natural compounds increase lysosomal enzyme activity (including cathepsins) and lysosomal membrane permeability (LMP) thereby leading to lysosome damage, release of lysosomal proteases and the so-called “lysosomal pathway of apoptosis” [45,46].

The importance of *SMARCA4* loss or overexpression during cancer initiation and progression has been debated, and the meaning for both modalities can be found in the literature [25,47]. The truncating mutations, fusions and homozygous deletion of *SMARCA4* co-occurred more frequently with *KRAS*, *STK1*1, and *KEAP1* mutations and are associated with shortest survival times of patients with non-small lung cancers (P < 0.001) [48]. On the contrary, analysis of the genomic data from the TCGA database of breast cancer patients showed <2% mutation frequency in invasive breast carcinomas, whereas the elevated expression of BRG1 occurred in 35-100% of analyzed primary tumors and was responsible for the high proliferation rate and was also a predictive biomarker for metastases risk [23]. However, the reversal of missense or truncating mutations has not been documented in the literature. The repeated exposure of a non-small lung cancer cell line with the *SMARCA4* truncating mutation caused a gain of missense mutation, which resulted in the inversion of the stop codon, restoration of mRNA translation and re-occurrence of BRG1. Since the initially truncating mutation occurred outside/beyond the enzyme functional domains, the short deletion in protein structure did not affect BRG1 activity or structure and the enzyme emerged functional, capable of controlling some ABC gene transcription and facilitating cell adaptation to paclitaxel in the environment. The transient deficiency of BRG1 in paclitaxel-resistant A549 cells sensitized these cells to a wide range of anticancer drugs. This implies the need to carefully testing SMARCA4 status, particularly when considering synthetic lethal therapy in SMARCA4 mutated cancers. Alteration in the BRG1 levels in cancer cells may drive their adaptation to particular conditions and promote transformation and progression of specific cancer types [49]. Furthermore, development of drug resistance is associated with considerable BRG1 redistribution in the genome of MDA-MB-231, which may support the gain of preferable transcriptome profile and desired phenotype. Although BRG1 was found at some ABCC promoters in non-resistant and resistant phenotypes, the activity of this enzyme varied. This implies the need for better understanding of the molecular mechanisms that determine gene transcription-dependence on BRG1, already difficult due to the multivalent DNA and chromatin-defined binding of BRG1 to histones as well as a relatively long list of SWI/SNF subunits and transcription co-factors that define BRG1 activity and interaction with particular gene promoters or enhancers [50].

In summary, pharmacological inhibition or degradation of SWI/SNF as well as BRG1 silencing leads to extralysosomal distribution of anticancer drugs, their deeper penetration of spheroids and substantial increase in drug cytotoxicity. Our study provides new target - SWI/SNF complex, and particularly SMARCA4 that encodes BRG1, for anticancer combinatorial interventions in paclitaxel-induced multidrug resistant phenotypes.

## Glossary

ABC: ATP-binding cassette
ADAADi: Active DNA-dependent ATPase A Domain inhibitor
CTX: cabazitaxel
DTX: docetaxel
LMP: lysosomal membrane permeability
MDR: multidrug resistance
PTX: paclitaxel
SRAP: Structurally Related Analog of PFI-3

## Acknowledgments

Graphical abstract was created with BioRender.com, agreement number: OF275Q4A5Q.

## Author contributions

Karolina Gronkowska: Investigation, Validation, Formal analysis, Writing – original draft. Sylwia Michlewska: Investigation, Visualization, Tomasz Płoszaj: Investigation, Data curation. Magdalena Strachowska: Investigation. Adrianna Stępień: Investigation. Maciej Borowiec: Resources, Writing – review & editing. Andrzej Bednarek: Resources, Writing – review & editing. Agnieszka Robaszkiewicz: Conceptualization, Validation, Investigation, Data curation, Resources, Funding acquisition, Project administration, Writing – original draft, Writing – review & editing.

## Data availability

Data underlying the results presented in this manuscript are provided within the Supplementary Data. Original raw data files containing sequence reads and quality scores were submitted to the NCBI Sequence Read Archive (SRA) database under the BioProject accession numbers: PRJNA1095909 and PRJNA1096152.

---

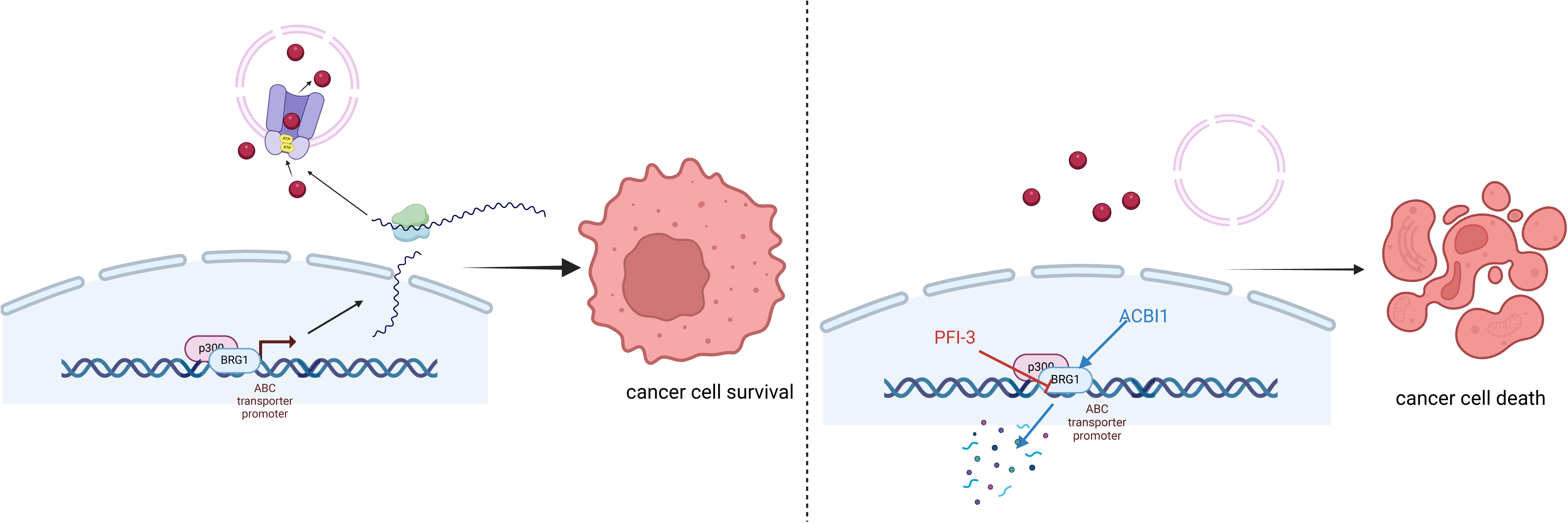

