## Supplementary figures for "BRG1 targeting overcomes ABCC-based multidrug resistance induced by paclitaxel"

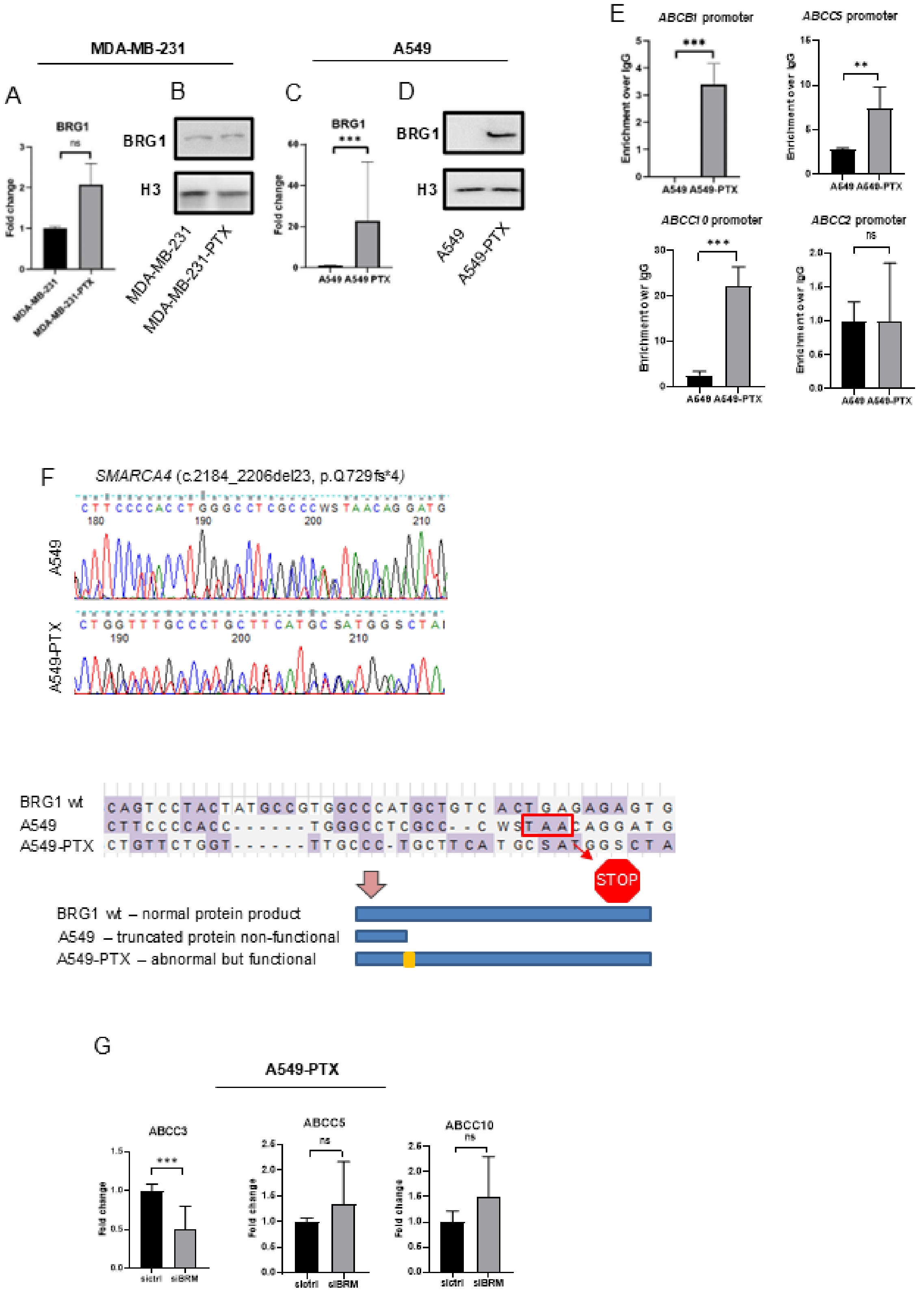


**Figure S1 Paclitaxel-induced changes in the nucleotide sequence of SMARCA4 led to reversal mutation that restored expression of functional BRG1 in paclitaxel-resistant A549 cells** (A, C) mRNA level of BRG1 was compared in non-resistant and ptx-resistant MDA-MB-231 (A) and A549 (C) cell lines by real-time PCR. Transcription level was normalized first to housekeeping genes (ACTB, GAPDH and HPRT1) and, then, mRNA level in basal cell lines was assumed as 1. The difference between two means was tested with Student's t-test, and statistically significant differences are marked with * when $p < 0.05$ *, ** when $p < 0.01$, *** when $p < 0.001$. (B, D) Protein level of BRG1 was compared in lysates of non-resistant and ptx-resistant MDA-MB-231 (B) and A549 (D) cells by western blot. Histone H3 was used as a loading control. (E) Recruitment of BRG1 to the promoters of *ABCB1*, *ABCC2*, *ABCC5* and *ABCC10* in two A549 phenotypes was monitored by ChIP-qPCR. Raw IP values were normalized first to input, then to the corresponding IgG controls. The difference between two means was tested with Student's t-test, and statistically significant differences are marked with * when $p < 0.05$ *, ** when $p < 0.01$, *** when $p < 0.001$ (F) Sanger sequencing results presented in Finch TV and aligned in MEGA11. (G) mRNA level of ABC transporters was compared between control and BRM-deficient samples by real-time PCR. Transcription level was normalized first to housekeeping genes (ACTB, GAPDH and HPRT1) and, then, mRNA level of control was assumed as 1. The difference between two means was tested with Student's t-test, and statistically significant differences are marked with * when $p < 0.05$ * and ** when $p < 0.01$.

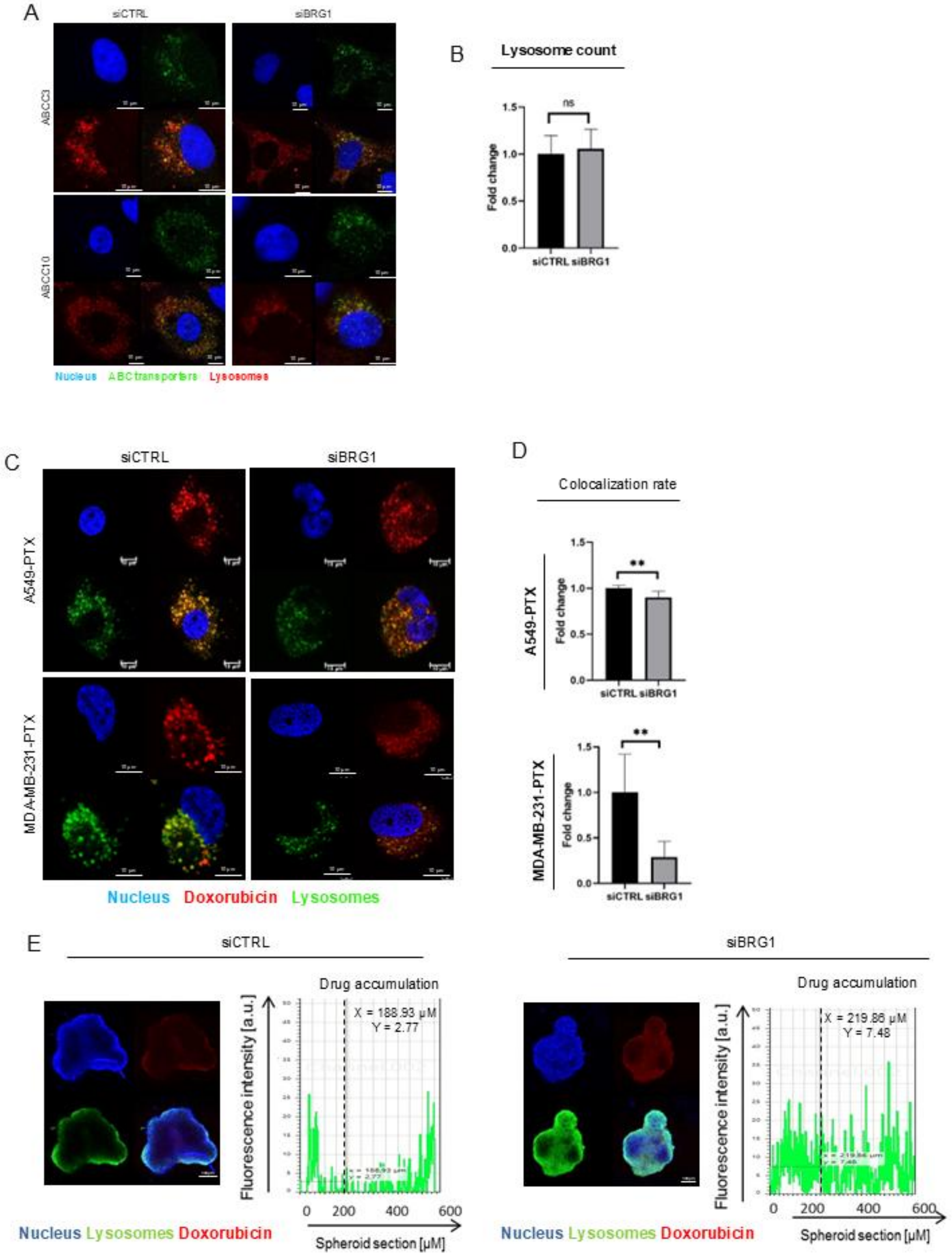


**Figure S2 BRG1 drives overexpression of ABC-transporters, which are enriched in lysosomes of PTX-resistant cell lines and crucial for reduced drug cytotoxicity due to drug sequestration in these organelles** (A) ABC transporter expression and localization 72 h after cell transfection with siCTRL and siBRG1 were visualized by immunocytostaining followed by confocal microscopy. Green fluorescence derived from Alexafluor488-conjugated secondary antibody corresponds to ABC transporters appearance in cells. DNA was stained with DAPI (blue). Lysosomes was stained with LysoTracker (red). (B) Lysosome count was compared between control cells and transiently silenced BRG1. The fluorescence intensity of lysosomal marker LysoTracker Deep Red was determined in arbitrary units (a.u.) with Leica Application Suite X. The difference between two means was tested with Student's t-test, and statistically significant differences are marked with * when $p < 0.05$ *, ** when $p < 0.01$, *** when $p < 0.001$ (C) BRG1 causes lysosomal sequestration of doxorubicin in paclitaxel-resistant cancer cell lines. Colocalization of doxorubicin and lysosomes was compared between BRG1 proficient and deficient cells. Autofluorescent doxorubicin is marked in red, lysosomes in green (LysoTracker Deep Red), DNA in blue (DAPI). The fluorescence intensity and colocalization of doxorubicin (D) and Paclitaxel Oregon Green (D) was determined in arbitrary units (a.u.) with Leica Application Suite X. The difference between two means was tested with Student's t-test, and statistically significant differences are marked with * when $p < 0.05$ *, ** when $p < 0.01$, *** when $p < 0.001$. (E) Doxorubicin accumulation in lysosomes of paclitaxel-resistant cells requires BRG1. Cells were grown for 3-weeks in 3D cultures. Control and BRG1 silenced paclitaxel-resistant A549 cell line spheroids were scanned for intracellular doxorubicin distribution. The autofluorescent drug is marked in red, DNA stained DAPI is blue, LysoTracker Deep Red stained lysosomes are green. The fluorescence intensity plot at spheroid cross section was created in arbitrary units (a.u.) with Leica Application Suite X.

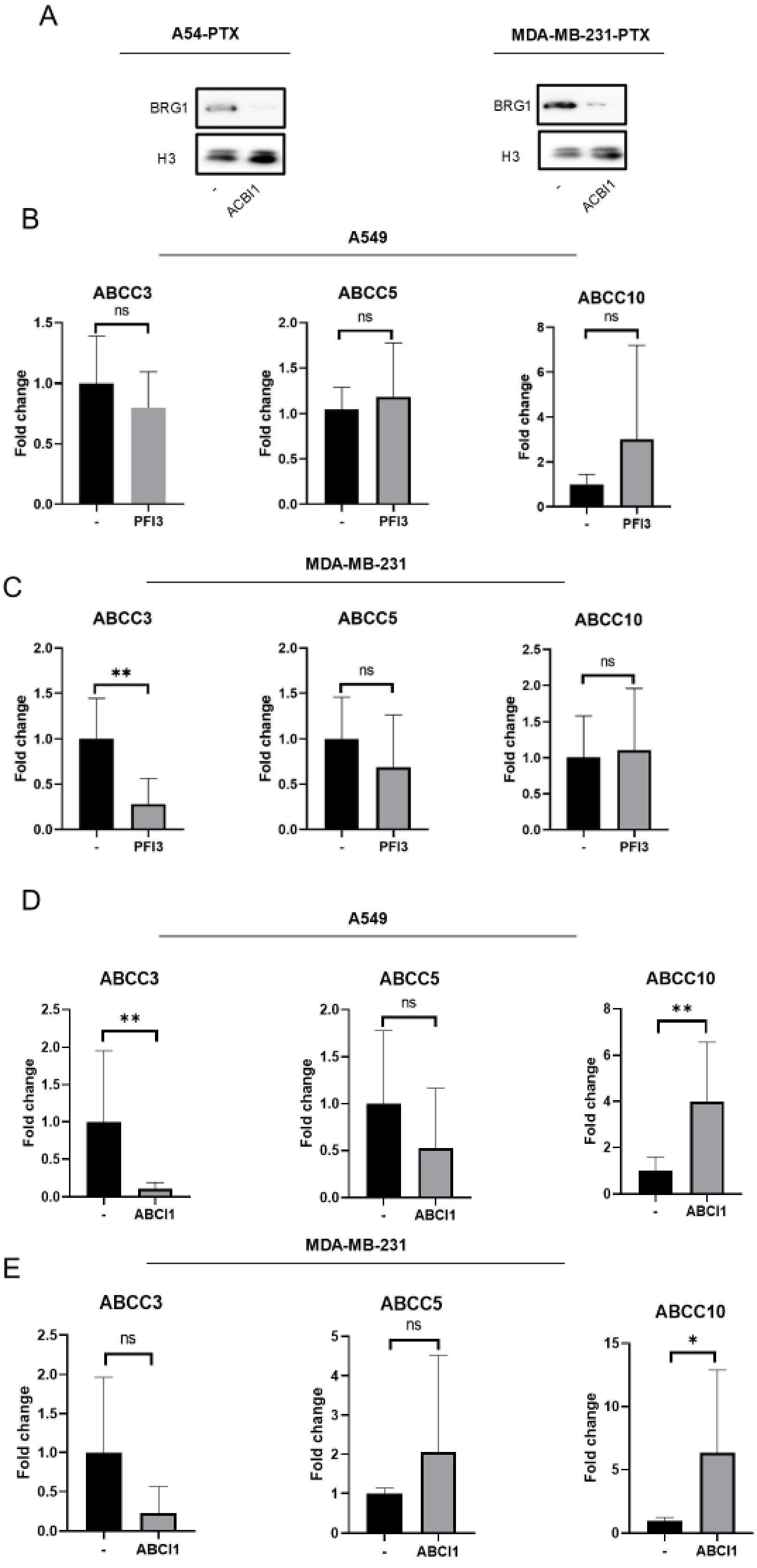


**Figure S3 SWI/SNF-targeting with PFI-3 and ACBI1 does not substantially reduces expression of ABCC5 and ABCC10 in A549 and MDA-MB-231 cells** (A)Western Blot representation of BRG1 degradation after ACBI1 treatment. H3 was used as a loading control. (B-E) The impact of SWI/SNF inhibitor – PFI3 (B,C) and SWI/SNF subunits degrader-ACBI1(D,E) on mRNA level of ABCC3, ABCC5 and ABCC10 was studied by real-time PCR. Cells were exposed to PFI3 at the concentration of 2.5 uM for 72h. Transcription level was normalized first to housekeeping genes (ACTB, GAPDH and HPRT1) and for control sample was assumed as 1. The difference between two means was tested with Student's t-test, and statistically significant differences are marked with * when $p < 0.05$ *, ** when $p < 0.01$, *** when $p < 0.001$.

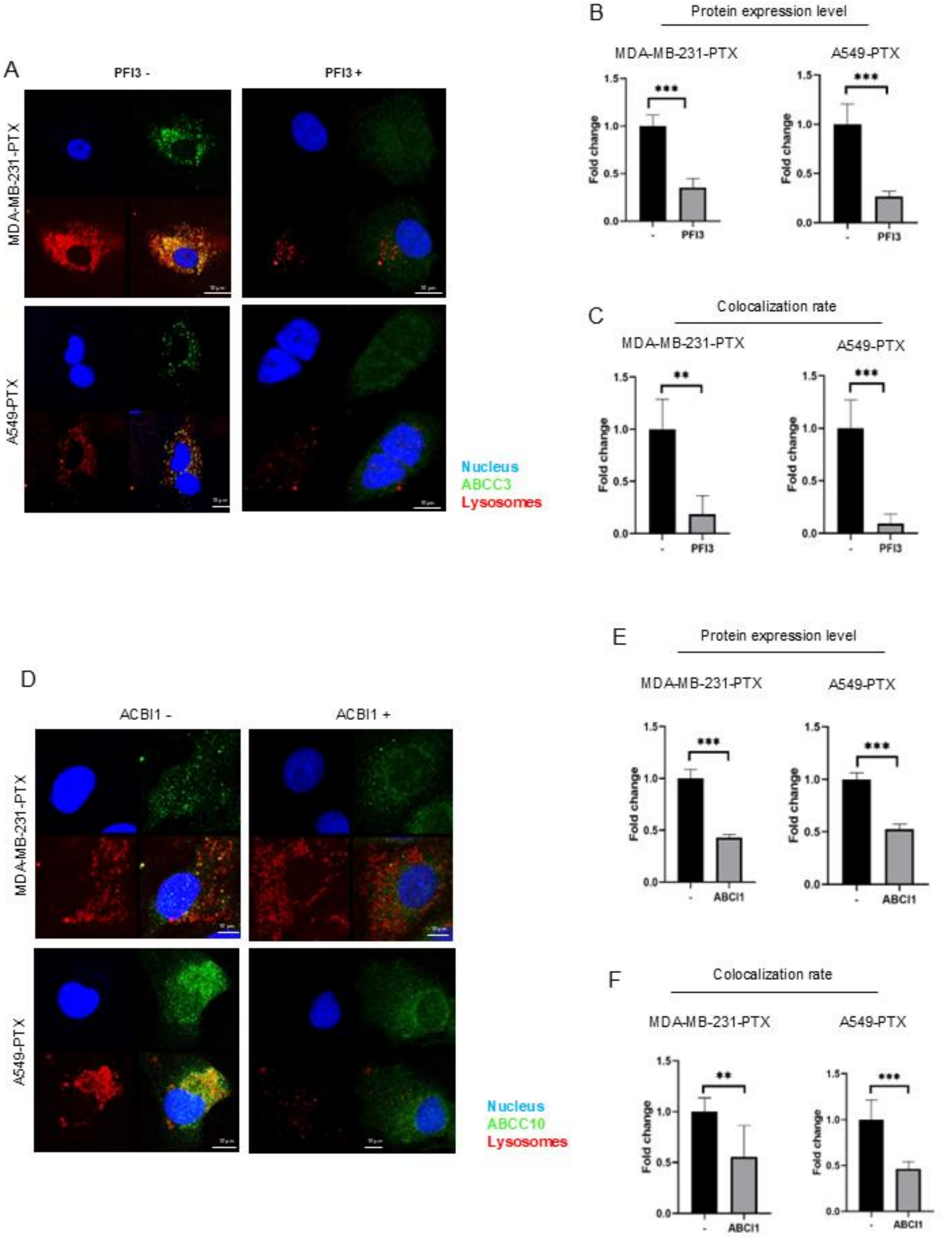


**Figure S4 SWI/SNF-targeting with PFI-3 and ACBI1 reduces expression of ABCC transporters overrepresented in lysosomes of paclitaxel-resistant cells** (A, D) ABC transporters expression and localization was visualized by immunocytostaining followed by confocal microscopy in non-treated vs PFI3 (A) and ACBI1 (D)-treated cells. Green fluorescence of Alexafluor488-conjugated secondary antibody represents ABCC transporters, blue fluorescence of DAPI - DNA, whereas lysosomal red fluorescence is from LysoTracker Deep Red. The fluorescence intensity (B, E) and colocalization (C,F) was determined in arbitrary units (a.u.) with Leica Application Suite X. The difference between two means was tested with Student's t-test, and statistically significant differences are marked with * when $p < 0.05$ *, ** when $p < 0.01$, *** when $p < 0.001$.

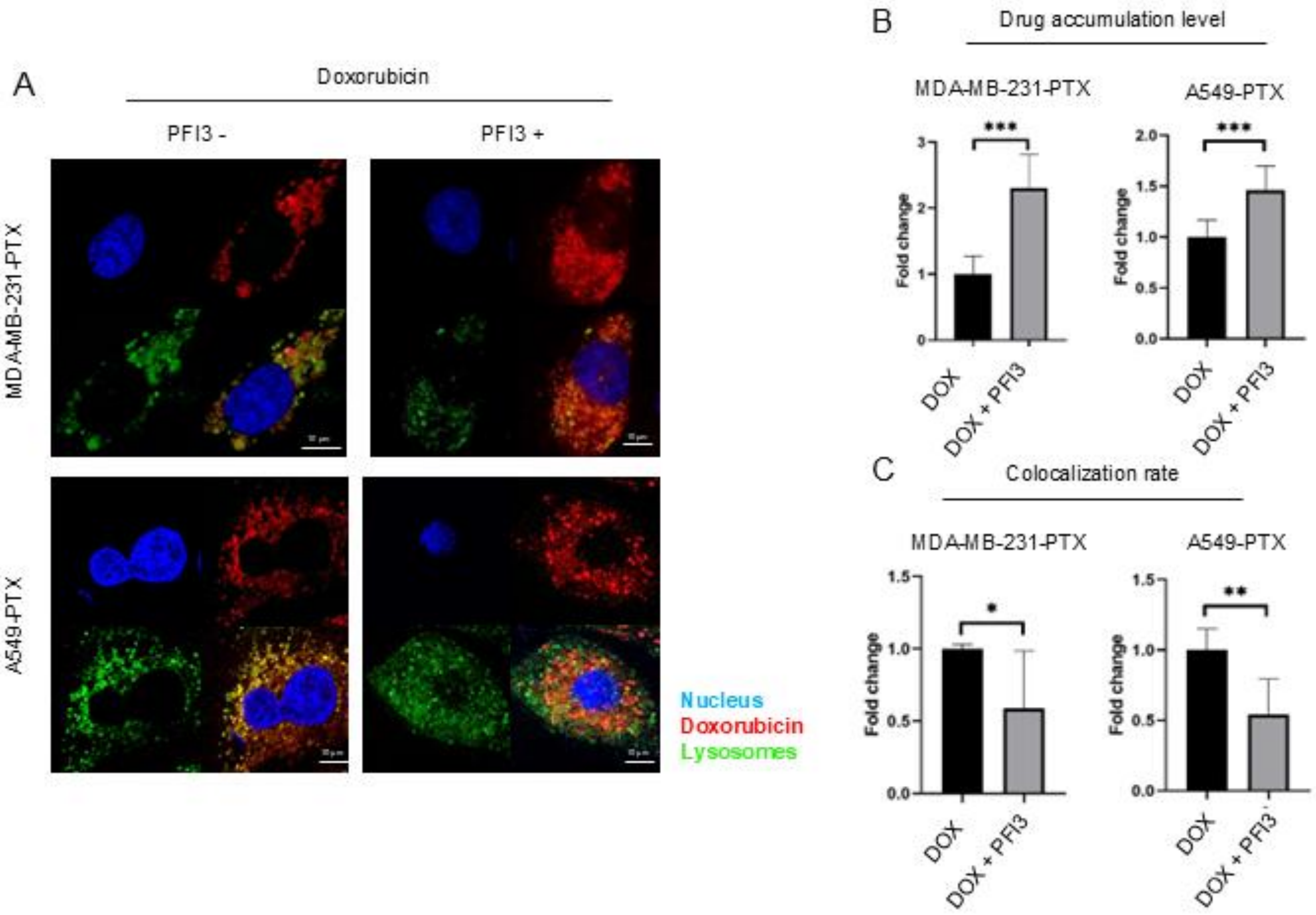


**Figure S5 Inhibition of SWI/SNF with PFI-3 decreases doxorubicin lysosomal uptake** (A-C) PFI3 uncouples doxorubicin accumulation in lysosomes of paclitaxel-resistant cells. (A) Intracellular localization of doxorubicin was analysed by confocal imaging in PFI3 treated (2.5 uM, 72h) and untreated cells. The autofluorescent drug is marked in red, DNA in blue (DAPI) and lysosomes in green (LysoTracker Deep Red). The fluorescence intensity of paclitaxel Oregon Green (B) and colocalization between the drug and lysosomes (C) was determined in arbitrary units (a.u.) with Leica Application Suite X. The difference between two means was tested with Student's t-test, and statistically significant differences are marked with * when $p < 0.05$ *, ** when $p < 0.01$, *** when $p < 0.001$
